# Structural Basis for Negative Regulation of ABA Signaling by ROP11 GTPase

**DOI:** 10.1101/2020.05.20.107185

**Authors:** Chuankai Zhao, Hassan Nadeem, Diwakar Shukla

## Abstract

Abscisic acid (ABA) is an essential plant hormone responsible for plant development and stress responses. Recent structural and biochemical studies have identified the key components involved in ABA signaling cascade, including PYR/PYL/RCAR receptors, protein phosphatases PP2C, and protein kinases SnRK2. The plant-specific, Roh-like (ROPs) small GTPases are negative regulators of ABA signal transduction by interacting with PP2C, which can shut off “leaky” ABA signal transduction caused by constitutive activity of monomeric PYR/PYL/RCAR receptors. However, the structural basis for negative regulation of ABA signaling by ROP GTPases remains elusive. In this study, we have utilized large-scale coarse-grained (10.05 milliseconds) and allatom molecular dynamics simulations and standard protein-protein binding free energy calculations to predict the complex structure of AtROP11 and phosphatase AtABI1. In addition, we have predicted the detailed complex association pathway and identified the critical residue pairs in AtROP11 and AtABI1 for complex stability. Overall, this study has established a powerful framework of using large-scale molecular simulations to predict unknown protein complex structures and suggested the molecular mechanism of the negative regulation of ABA signal transduction by small GTPases.

## 1 Introduction

The plant hormone abscisic acid (ABA) regulates a variety of developmental processes and responses to environmental stresses in plants.^1–4^ Recently, the core components involved in ABA signaling network have been identified, including the family of PYR/PYL/RCAR (PYLs, pyrabactin resistance 1/PYR1-like/regulatory component of ABA receptor) ABA receptors, protein phosphatase PP2Cs (clade A serine/threonine protein phosphatase 2C) and protein kinase SnRK2s (subfamily 3 SNF1-related kinase 2).^5^ Under non-stress conditions, PP2Cs bind to SnRK2s and dephosphorylate SnRK2s, resulting in the inactivation of SnRK2s (Fig. 1A).^6,7^ Under stress conditions, plants promote *in planta* synthesis of ABA molecules and trigger the negative regulatory ABA signaling network.^8^ When ABA binds to PYLs, PYLs undergo pronounced conformational change to facilitate PYL-PP2C interactions via direct binding.^9–12^ Upon being free from inhibition by PP2Cs, SnRK2s activate through autophosphorylation, and then phosphorylate downstream signaling components, eventually triggering a range of ABA responses (Fig. 1B).^13–15^ In *Arabidopsis thaliana*, there are 14 functionally redundant PYLs. PYL4-13 exist as a monomer under physiological conditions^10,11,16,17^ and can bind to and inhibit PP2Cs in the absence of ABA, which would theoretically cause “leaky” ABA signal transduction in the absence of stimulus. ^16,18^

**Figure 1:**
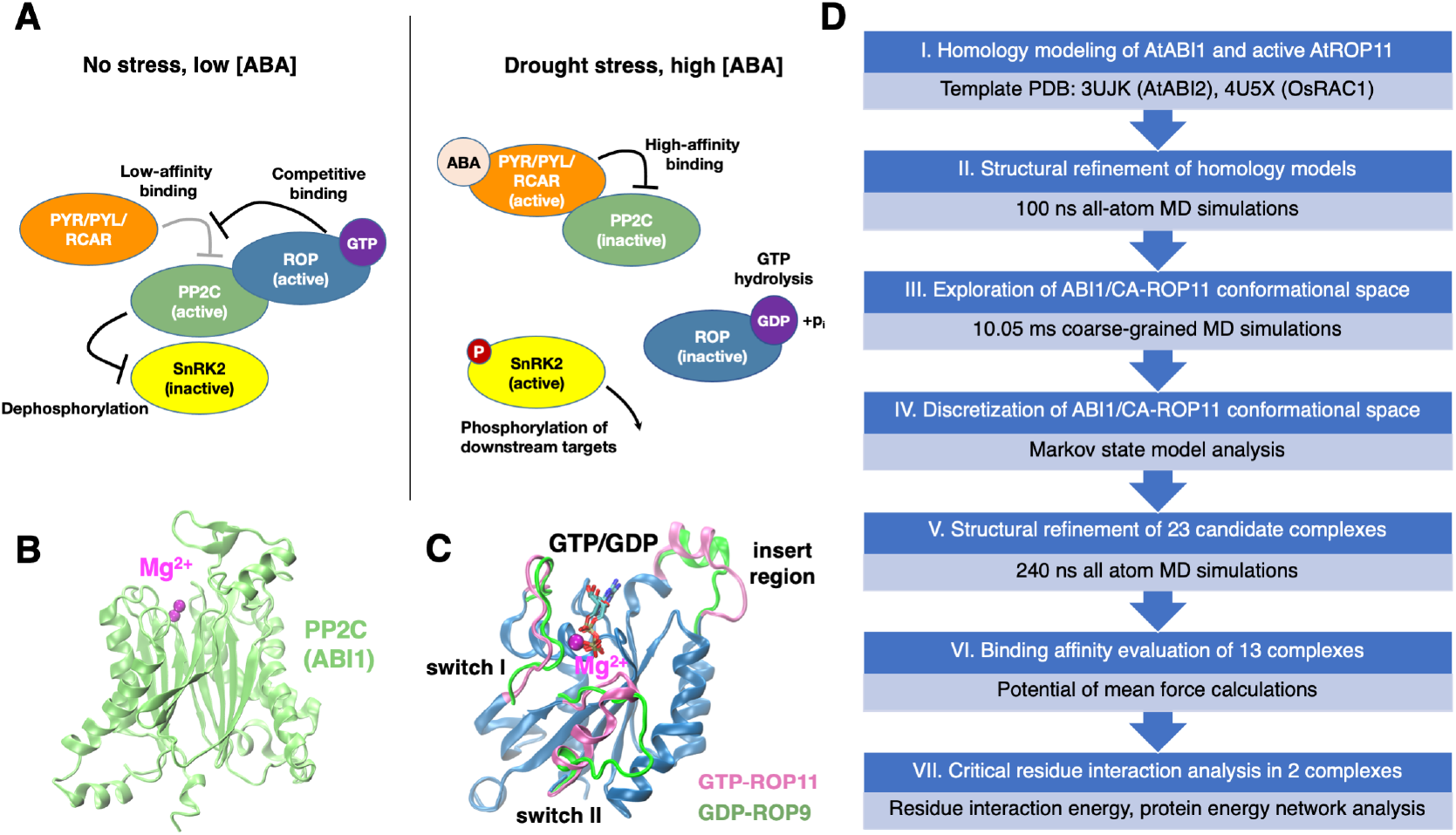
Schematic of negative regulation of ABA signaling by ROP GTPase and atomic structures of ABI1 and ROP11. (A) Under normal conditions, ROP GTPase competitively binds to PP2C and prevents ABA-independent inhibition of PP2C, leading to no stress responses. Under stress conditions, ABA binds to PYR/PYL/RCAR receptors and results in high-affinity binding between ABA-bound receptor and PP2C, triggering downstream ABA responses. Molecular structures of (B) ABI1 and (C) GTP-bound active ROP11 (homology model from PDB ID: 4U5X) and GDP-bound inactive ROP9 GTPase (PDB ID: 2J0V^24^). The three regions in ROP11, including insert region and switch I and II, that undergo major conformational changes upon activation are highlighted in pink (active) and green (inactive). (D) Overview of the computational workflow for predicting ABI1-ROP11 complex structure.

Genetic and biochemical studies have identified the plant-specific Rho-like (ROPs) small GTPases as negative regulators of ABA signaling, which can potentially shut off ABA signaling in the absence of stimulus.^19–23^ Li *et al*. have demonstrated that expression of a constitutively active ROP11 (CA-ROP11, Q66L), a member in a family of 11 ROPs, can suppress a variety of ABA-mediated responses, including seed germination, seedling growth, stomatal closure and plant responses to drought stress.^20^ Furthermore, Li *et al*. have shown that ROP11 negatively regulates ABA signal transduction by stabilizing the activity of ABI1 (a member of PP2C family, Fig. 1B).^20,23^ Based on these results, they have proposed a model that can effectively shut off ABA signal transduction in the absence of ABA (Fig. 1A).^23^ In this model, ROPs serve as signaling switches by adopting either a GDP-bound inactive state or a GTP-bound active state (Fig. 1C). Under non-stress conditions, monomeric PYLs bind to PP2C in the absence of ABA with a low affinity. GTP-bound ROPs can competitively bind to PP2C and interfere with the leaky repression of PP2C activity by monomeric PYLs (Fig. 1A).^20,21,23^ Under stress conditions, ABA-bound PYLs bind to PP2Cs with an enhanced affinity, which would put ROPs at a competitive disadvantage. Meanwhile, ABA leads to the inactivation of ROPs by adopting the GDP-bound state.^23^ As a result, in the presence of ABA, PP2Cs are fully inactivated by PYLs, releasing SnRK2s to induce ABA responses (Fig. 1B). Overall, ROPs, notably ROP10 and ROP11, play a critical role in regulating ABA signal transduction, while molecular understanding of how ROPs competitively bind to PP2Cs and stabilize PP2C activity is still lacking.

In recent years, a plethora of molecular modeling techniques have emerged as powerful computational tools for structural prediction and binding affinity evaluation of protein complexes, including docking^25–27^ and molecular dynamics (MD).^28–30^ Docking is useful for predicting the molecular assembly of proteins given their individual structures, while even the state-of-the-art docking techniques are limited in incorporating protein dynamics and cofactor-protein interactions. MD simulations can capture the motion of proteins and their associated cofactors with atomistic details.^31,32^ With the rapid advances of computing power, long timescale MD simulations have been routinely used in studying protein dynamics and function, ^33–35^ such as protein conformational change,^36,37^ protein-ligand binding^38–42^ and protein-protein association. ^29,30^ We have recently utilized large-scale all-atom MD simulations to study the molecular mechanism of ABA-mediated activation of three sub-type ABA receptors^43,44^ and predict a near-native complex structure of ABA receptor PYR1.^45^ In contrast to all-atom MD simulations, coarse-grained (CG) MD simulations are useful in addressing the challenges of simulating long-timescale protein dynamics,^46,47^ such as protein-protein association.^47–49^ Generally, proteins are represented with beads and simulated with less details in CG MD simulations, where each bead approximates multiple heavy atoms and their connected hydrogen atoms. To model protein complex structure, one can perform large-scale CG MD simulations to sample the vast space of complex configurations, and subsequently utilize binding affinity evaluation to rank candidate complexes from simulations. ^47–49^

In this study, we have performed extensive MD simulations to predict the complex structure of ROP11 and ABI1 in *Arabidopsis thaliana* (denoted as AtROP11 and AtABI1) (Fig. 1D, Table S1). Since there is no GTP-bound and active structure available for ROP11, we have predicted the structure using homology modeling based on the crystal tructure of active OsRac1 in rice (with 82.95% sequence identity to AtROP11). In order to explore the conformational space of ROP11-ABI1, we performed 10.05 ms adaptive CG MD simulations to sample the complex structural ensemble of ABI1 and a constitutively active ROP11 (CAROP11). We analyzed the large-scale CG MD simulation data using a statistical approach called as Markov state model, ^35,50^ which was used to discretize the ensemble into 600 individual states along with quantitative estimation of their equilibrium populations. We chose the top 25 candidate complex structures that account for more than 80% of total population, and obtained their atomic structures by aligning the atomic structures of ROP11 and ABI1 to these complexes. Next, we performed 240 ns all-atom MD simulations to further refine these complex structures, resulting in 13 complexes that were stable. To further differentiate the 13 complexes, we performed potential of mean force (PMF)-based energy calculations to evaluate the relative stability of these complexes. We obtained 2 candidate structures which demonstrate relative high stability. Based on the analysis of residue interaction energies in the predicted complexes, we identified and characterized the critical residue pairs responsible for forming the complexes. Overall, this study has elucidated the structural basis for negative regulation of ABA signaling by ROP GTPase, which can create new avenues for engineering ABA signaling network to control ABA-mediated responses in plants.

## 2 Results

### Homology models of GTP-bound, active ROP11 and ABI1

The GTP-bound, active AtROP11 structure was predicted from homology modeling using the active OsRac1 in rice (PDB ID: 4U5X^51^) as the structural template. The crystal structures of several active GTPase with certain degrees of sequence variation (currently available in Protein Data Bank) suggested that the active structure of GTPase is conserved (Fig. S1), which justifies the accuracy of our homology model of AtROP11. The modeled structure was further refined by running 100 ns all-atom MD simulations. Compared to the crystal structure of GDP-bound and inactive AtROP9, the major conformational changes are observed in three regions (Fig. 1C), named as switch I, switch II and insert region, which are known to be critical for small GTPase activation. ^51^ Notably, the helix in switch II of the active structure unfolds in the inactive state. The conformational changes in switch I and insert region are less pronounced. The conformational changes in ROP11 upon activation may be critical for protein-protein interaction between ROP11 and ABI1. Based on this homology model, an atomic structural model of the constitutively active ROP11 (CA-ROP11, Q66L) ^20,21^ was generated by performing *in silico* side chain mutation. ABI1 structure was predicted from the free crystal structure of its homolog ABI2 (PDB ID: 3UJK^7^), which only deviates from the crystal structure of ABI1 (in complex with ABA receptor, PDB ID: 3JRQ ^52^) by 1.42 Å (Fig. S1).

### 2.1 Extensive CG MD simulations capture the conformational space of ABI1/CA-ROP11 complex

We sought to perform large-scale explicit solvent CG MD simulations using available ABI1 and CA-ROP11 structures, in order to sample possible configurations of ABI1 and ROP11 complex. We simulated CA-ROP11 to mimic the interactions between ABI1 and GTP-bound, active ROP11. Martini coarse-grained force field for proteins was used in our CG MD simulations, which has been used extensively in mesoscale modeling of biomolecule and soft matter.^53,54^ The atomic structures of ABI1 and CA-ROP11 were converted to the bead representation using the four-to-one mapping strategy required by the Martini force field (Fig. S2). Generally, the heavy atoms in the backbone of each amino acid are mapped to one bead, and the heavy atoms in the side chain of each amino acid are mapped to 1 or more beads of various types (e.g. polar, non-polar, apolar, and charged). In total, 5 rounds of CG MD simulations were performed, resulting in 10.05 ms aggregate simulations including 950 independent trajectories of 10 or 21 *µ*s each (supplementary methods, Table S2). We have observed that ABI1 and CA-ROP11 mostly reached a relatively stable configuration within 10 or 21 *µ*s of each trajectory.

In order to identify the near-native complex from the coarse-grained conformational ensemble of ABI1 and CA-ROP11, we then utilized Markov state models (MSMs) to analyze the simulation data. MSMs is a powerful tool for analyzing large-scale MD simulation data on protein dynamics.^35,50^ It characterizes protein dynamics by discretizing entire protein conformational space into a certain number of states and the inter-state transition probabilities between these states.^35,50^ The discretization step is usually achieved by clustering the conformations according to their structural similarities and kinetics. The transition probabilities between different clusters are estimated statistically based on the inter-cluster transitions captured from CG MD trajectories. The equilibrium populations of individual clusters can then be estimated from the transition probability matrix of MSMs, which serve as a metric to rank the likelihood of each cluster as the native complex. In our study, we have defined six metrics for clustering ABI1 and CA-ROP11 conformations, including the center-of-mass distance between two proteins and five angles or dihedral angles that measure the relative orientation and position of two proteins (Fig. S3). We have further employed time-lagged independent component analysis (tICA)^55^ on the six metrics to generate the slowest relaxing degree of freedoms (denoted as tICs) in ABI1/CA-ROP11 association processes. We then clustered all the snapshots into 600 states based on 4 tICs, and an MSM was constructed using a lag time of 800 ns (supplementary methods, Fig. S4).

Using the MSM, we obtained 600 representative configurations of ABI1/CA-ROP11 along with the probabilities of observing these states at equilibrium. We have also generated the conformational free energy landscape to characterize complex thermodynamic stability (Fig. S5A,B). The minima on the landscape corresponding to relative stable complex conformations (Fig. S5A,B). We then focused our further analysis on the top 25 states with the highest equilibrium populations. The sum of the populations of these 25 states is greater than 80% of total populations (Fig. S5C). In addition, the 25 states cover all the minima of the free energy landscape (Fig. S5A,B), suggesting that the native complex is likely among these 25 states. Overall, large-scale CG MD simulations and MSM analysis have generated the top 25 candidate complex structures of ABI1 and CA-ROP11.

### 2.2 Top 13 complexes obtained after structural refinement by allatom MD simulations

We then converted the CG models of top 25 states into atomic representations, which was achieved by inverse one-to-four mapping followed by short MD simulations to equilibrate the atomic structures (supplementary methods). Due to the possible inaccuracy of inverse mapping, we have aligned the initial atomic structures of ABI1 and GTP-bound, active ROP11 to the converted atomic complex structures. In this way, we have obtained 25 candidate complex structures with GTP bound to ROP11. By visually inspecting these structures, we observed that ABI1 and ROP11 in three states (top 3, 7 and 11 states in Fig. S5B) share similar conformations. For further analysis, we combined the three states as a single state and also summed their equilibrium populations.

To further refine the interfacial interactions for the 23 candidate complexes, we have performed two independent all-atom MD simulations starting from each complex structure for 240 ns. We have characterized the center-of-mass distance *r* between two proteins and the root mean square deviation (RMSD) of complex from initial structure with respect to simulation time (Figs. S6 and S7). For each of these states, if large fluctuation of either *r* or RMSD (*>*5 Å) is observed in either trajectory, it suggests that the complex structure is not stable in all-atom MD simulations. Using this criteria, we ruled out 10 candidate complexes, leading to 13 remaining candidate complexes as shown in Fig. 2. Among the 13 states, the binding of ROP11 in several states would exclude the binding of ABA receptor to ABI1, whereas, in the other states, the binding site of ROP11 does not significantly overlap with the binding site of ABA receptor (Fig. 2). Overall, extensive all-atom MD simulations have further refined the structural models from CG MD simulations and helped downselect the 13 complex structures from 23 candidate complex structures.

**Figure 2:**
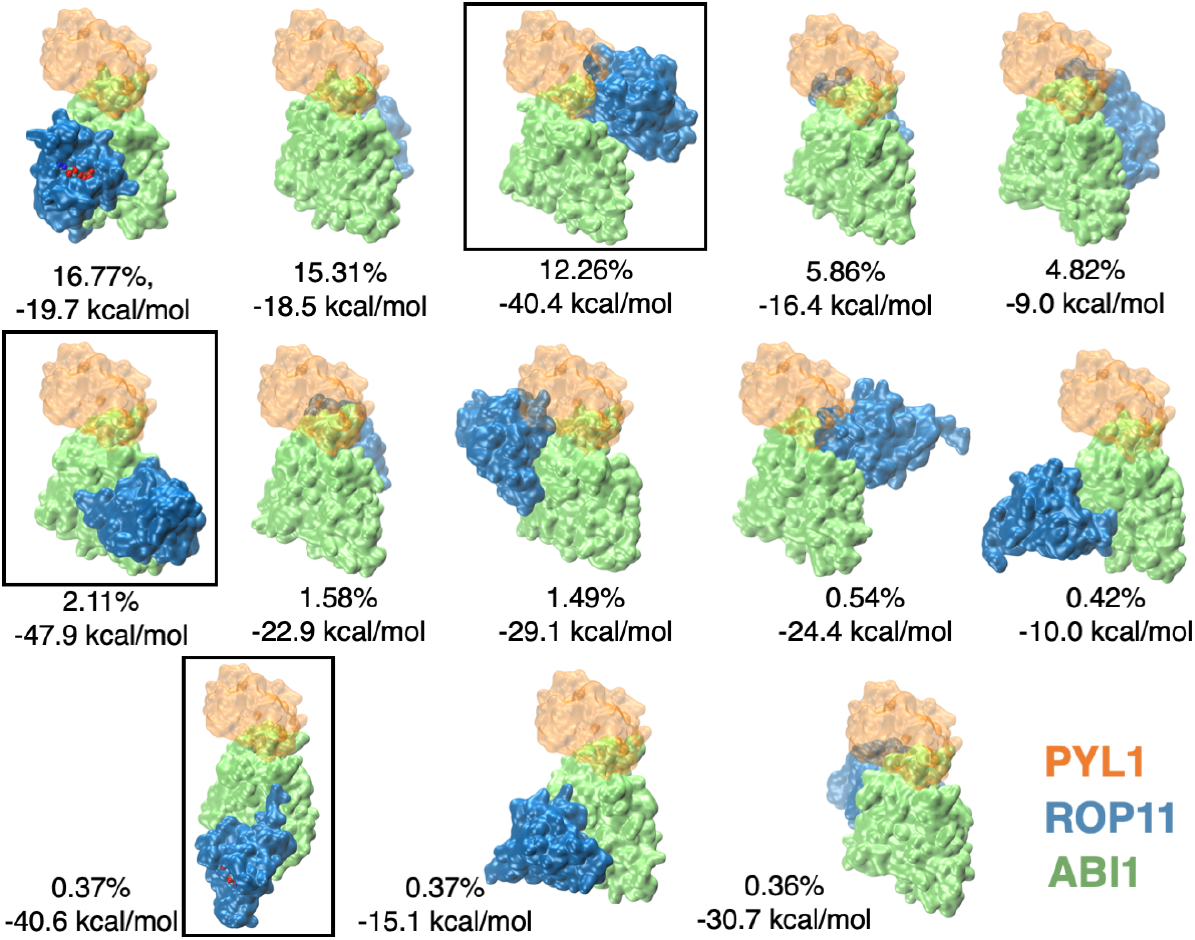
Snapshots of the top 13 candidate complex structures after structure refinement by all atom MD simulations. ABI1 and ROP11 are shown in lime and blue, respectively. ABA receptor PYL1 (orange) is also shown to indicate PYL1-ABI1 binding site. The equilibrium populations and the PMF depths for these complex structures are shown. The three states highlighted in black boxes are state 160, state 240, and state 3, as named according to their cluster numbers in the MSM.

### 2.3 Potential of mean force calculations for quantitative characterization of the relative stability of candidate complexes

In order to further differentiate the 13 atomic structural models of ABI1 and ROP11, we sought to perform free energy calculations to evaluate the thermodynamic stability of 13 complexes. In this part, we utilized potential of mean force (PMF) calculations to evaluate the free energy profile to separate each candidate complex structure from bound state to a fully unbound state (supplementary methods). ^56–58^ By checking the PMF depth between the bound state and the unbound state, we can quantitatively characterize the relative stability of these complexes. For each complex, we separated ABI1 and ROP11 along a vector *r*, which connects the center of mass of two proteins, in the presence of geometrical restraints (Fig. 3A). To determine the separation PMF, we selected a series of complex conformations with *r* evenly distributed in a certain range from the associated state to the fully dissociated state. Replica-exchange umbrella sampling (REUS) MD simulations were started from these complex conformations, with a harmonic potential acting on *r* to restrain the distance between ABI1 and ROP11 (Table S3). From these simulations, the separation PMF can then be estimated using the statistical free energy method, multistate Bennett acceptance ratio (MBAR).^59^ The purpose of applying these geometrical restraints was to accelerate the convergence of separation PMF, including the conformational restraints (*B*_*ABI*1_, *B*_*ROP* 11_) and the restraints on the relative position (Θ, Φ) and orientation (*ψ, ϕ, θ*) of two proteins (Fig. 3A). Using this protocol, we have obtained the separation PMFs for the 13 complexes (Fig. S8) and the PMF depths are indicated in Fig. 2.

**Figure 3:**
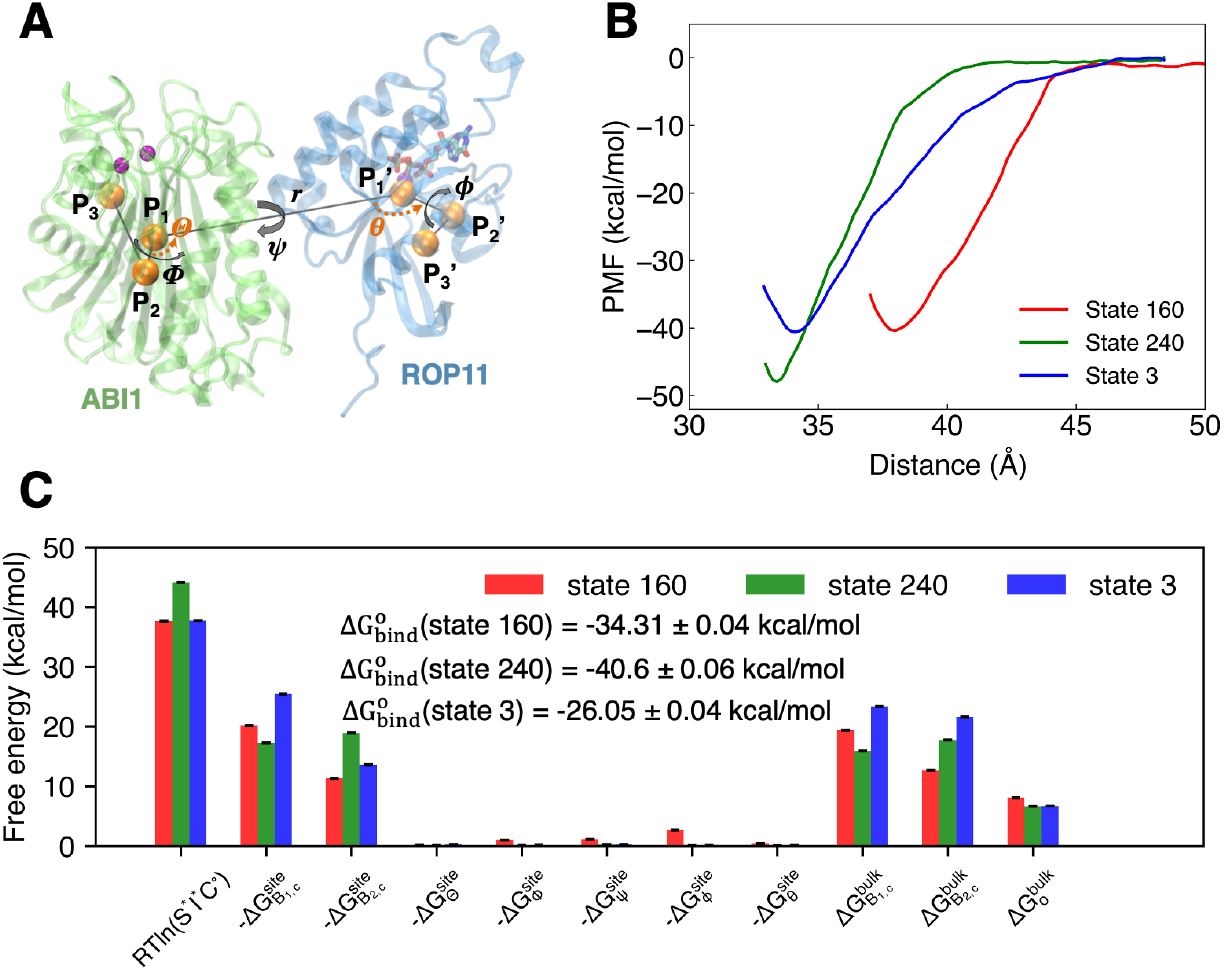
Determination of standard binding free energies for three candidate complex structures. (A) Snapshot of ABI1 and ROP11 and the collective variables used in separation REUS MD simulations. The center of mass distance between ABI1 and ROP11 (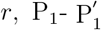 distance), and the Euler angles 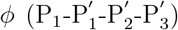 and 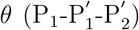 defines the relative position between two proteins. The Euler angles, 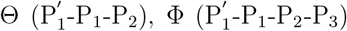, and 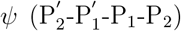, relate the relative orientation between the two protomers. In addition, the conformations of ABI1 and ROP11 are restrainted by RMSDs of the two proteins (*B*_*ABI*1_, *B*_*ROP* 11_) from the initial strutcures. (B) Potential of mean force (PMF) profiles for the separations of ABI1 and ROP11. The error bars on the PMFs are shown. (C) Free energies associated to the components of 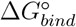 for three different complexes. The error bars for all components are less than 0.04 kcal/mol.

From the PMF depths, we have identified 3 states (third, sixth and eleventh states in Fig. 2, denoted as state 160, state 240, state 3 according to their cluster numbers in the MSM) with PMF depths greater than 40 kcal/mol (Fig. 3B), which indicates their higher stability as compared to the other 10 complexes. This suggests that the three candidate complexes are more likely to be the native complex of ABI1 and ROP11. We further sought to determine standard protein-protein binding free energy (Δ*G*^*o*^) for the three candidate complexes (supplementary methods). Δ*G*^*o*^ includes the contribution from the integration of separation PMF and the contributions from the applied conformational and angular restraints during separation PMF calculations. The contribution of separation PMF to Δ*G*^*o*^ (*RTln*(*S*^∗^*I*^∗^*C*^*o*^), including the contributions of applied restraints) can be calculated by numerical integration of separation PMF. To obtain Δ*G*^*o*^, the contributions of applying the restraints in the bound state and removing them in the unbound states should also be calculated. We thus have determined a series of PMFs relating to each individual restraints (Fig. S9) and calculated their contributions to Δ*G*^*o*^ (Fig. 3C). Take altogether, Δ*G*^*o*^ for state 160, state 240 and state 3 are −34.31 ± 0.04 kcal/mol, −40.6 ± 0.06 kcal/mol, and −26.05 ± 0.04 kcal/mol. Overall, these free energy calculations suggest that state 160 and state 240 are the most probable complexes of ABI1 and ROP11.

### 2.4 Structural analysis of the candidate complexes predicted from MD simulations

We further sought to examine the three candidate complex structures and see if each of these structures agree with known experimental data. Due to the nature of competitive binding between ABA receptor and ROP11 to ABI1,^20–23^ the binding of one to ABI1 would theoretically exclude the binding of the other to ABI1 and therefore their binding site would likely partially overlap. However, the binding of ROP11 should not block the catalytic site of ABI1 since ROP11 stabilizes ABI1 activity. ^20–23^ In addition, only GTP-bound and active ROP11 can interact with ABI1 and the conformational changes in ROP11 upon activation are observed in switch I, switch II and insert region, suggesting that the three regions are likely to be involved in the interface.

As shown in Fig. 2, the binding site of ROP11 in the state 160 (the third snapshot in Fig. 2) has partial overlap with the binding site of ABA receptor PYL1. In PYL1-ABI1 structure, W300 is docked into the binding pocket of ABA receptor, whereas W300 is involved in forming the interface between ABI1 and ROP11 (Fig. 4A,B). These results suggest that the binding of ROP11 in the state 160 would exclude the binding of PYL1 to ABI1, which is consistent with the previous experimental results. ^20–23^ ROP11 adopts a significantly different binding pose compared to PYL1, which leaves the catalytic site exposed to solvent. This is consistent with the fact that ROP11 stabilizes ABI1 activity. ^20–23^ The switch I in ROP11 interacts with the catalytic site of ABI1, which is similar to that the activation loop of SnRK2 interacts with the catalytic site in ABI1 (Fig. 4B,C). The key interaction involved at the interface is through electrostatic interaction between Mg^2+^ in the catalytic site of ABI1 and D36 in switch I of ROP11 (Fig. 4D,E). In addition, K32 and K163 in ROP11 forms electrostatic interaction with D278, D351 and D282 near the catalytic site of ABI1. N301 in ABI1 forms polar interaction with GTP and polar interactions are formed between W300-D127, Q408-N44, and R409-T29 in ABI1 and ROP11. The bulky side chain of W300 stabilizes the hydrophobic interaction between the loop in ABI1 and insert region of ROP11. We have performed ~520 ns MD simulation on the state 160, and the complex remains highly stable within the simulation timescale (Fig. S10). Finally, the GTP binding site of ROP11 in the state 160 remains largely exposed, allowing for GTP/GDP exchange catalyzed by guanine-nucleotide exchange factor (GEF) enzyme. ^22,23^ We identified the association pathway between ABI1 and CA-ROP11 as captured in our CG MD simulations (Fig. 5). The insert region in CA-ROP11 initially recognizes the loop in ABI1 through electrostatic interactions between the charged residue pairs in ABI1 and ROP11. CA-ROP11 then adapts its orientation to finally assume the binding pose in the state 160. The association pathway also highlighted the essential role of switch I and insert region of ROP11 in forming the complex. Overall, this candidate complex structure has relatively large equilibrium population and separation PMF depth, and agrees with the competitive binding between ROP11 and ABA receptor as well as the activation process of ROP11.

**Figure 4:**
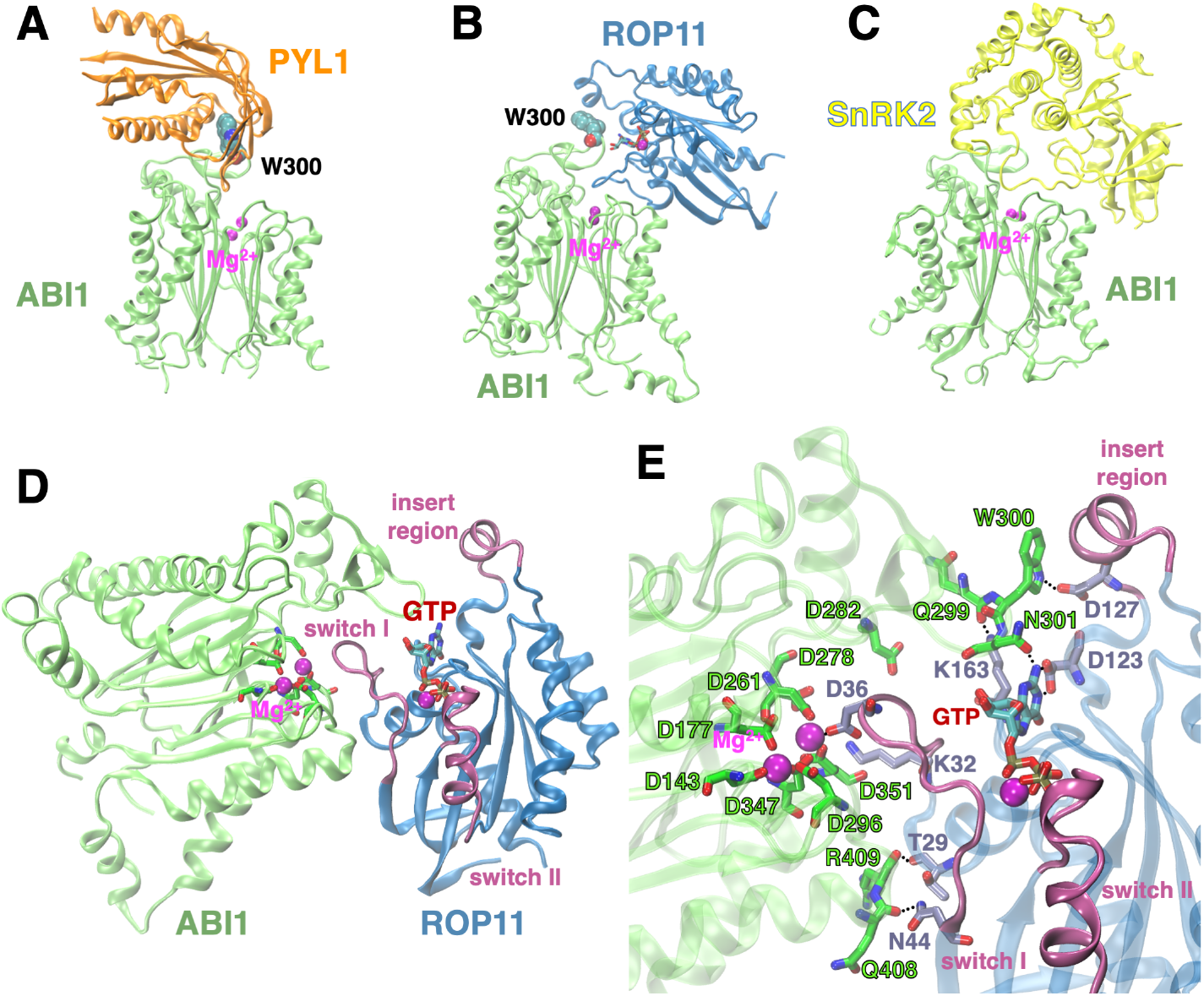
ABI1-ROP11 candidate complex structure (state 160). Molecular structures of (A) PYL1 and ABI1 (PDB ID: 3JRQ^52^), (B) ROP11 and ABI1 (predicted), and (C) SnRK2 and ABI1 (homology model from PDB ID: 3UJG^7^). (D) ROP11 GTPase binds to ABI1 through the switch I region, and leaves the catalytic site of ABI1 and GTP binding site of ROP11 exposed to solvent. (E) D36 in ROP11 GTPase interacts with Mg^2+^ at the catalytic site of ABI1, promoting ABI1-ROP11 interactions. Polar interactions between ABI1 and GTP/ROP11 are highlighed.

**Figure 5:**
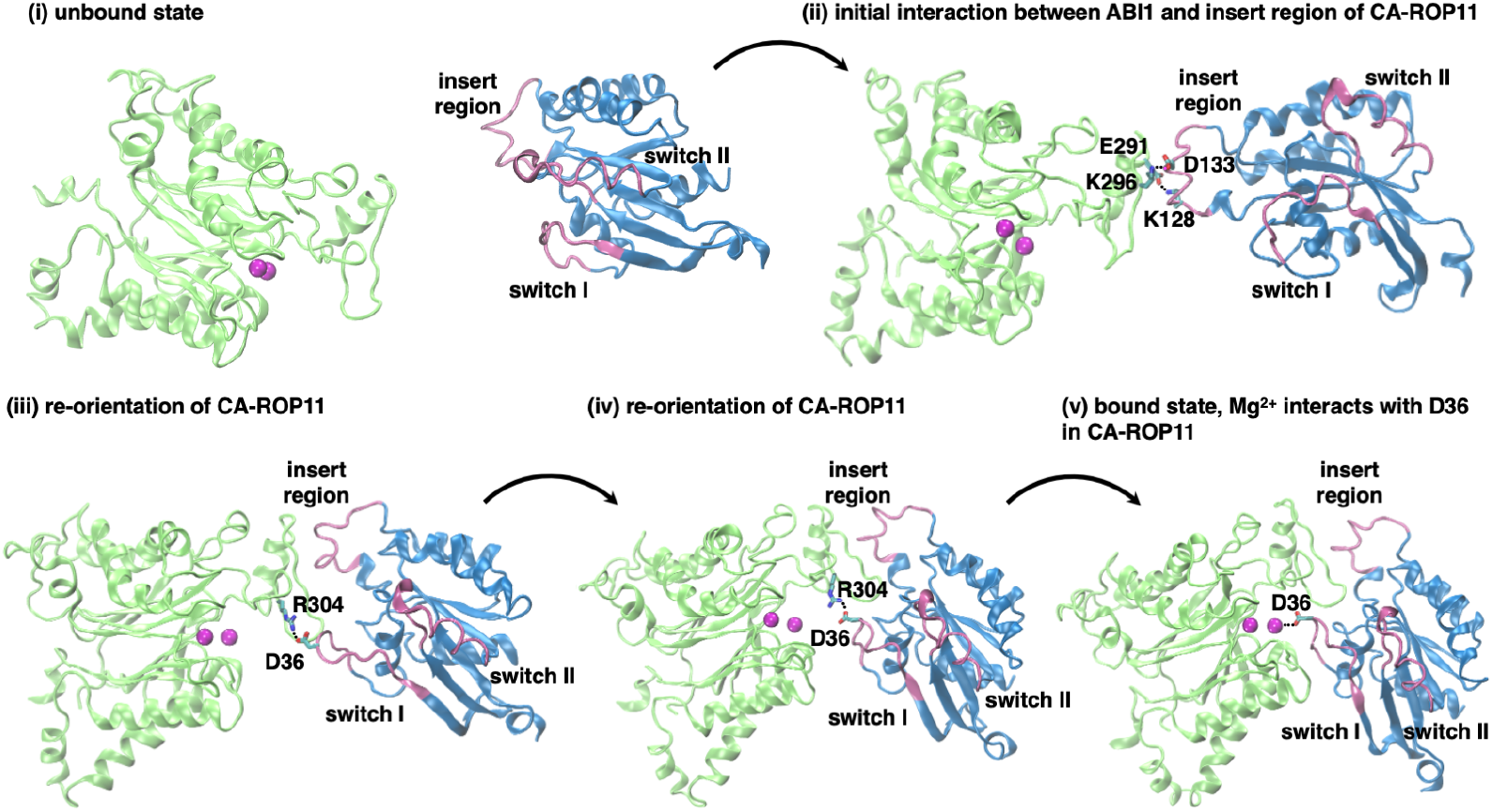
The association pathway between ABI1 and CA-ROP11 to form state 160 predicted from CG MD simulations. The initial interaction between ABI1 and CA-ROP11 is mediated by the charged residues (E291-K128, K296-D133) in ABI1 and the insert region of CA-ROP11. Next, CA-ROP11 adapts the orientation to rotate around ABI1, facilitated by the interaction between R304 in ABI1 and D36 in the switch I of CA-ROP11. Finally, D36 recognizes the Mg^2+^ in the catalytic site of ABI1, and CA-ROP11 binds to ABI1. This pathway was captured in continuous trajectory from CG MD simulations. The atomic complex structures were converted from the snapshots in CG MD trajectory.

For both state 240 and state 3, the binding of ROP11 would not directly impact the binding of PYL1 to ABI1 as shown in Fig. 2 (the sixth and the eleventh snapshots). In the state 240, the GTP is directly involved at the interface between ABI1 and ROP11 (Fig. S11A). Also, the switch I, switch II and insert region in ROP11 are directly involved in the interaction between ABI1 and ROP11 (Fig. S11A). At the interface, there are several charged residue pairs (R189-D127, E190-K128 and K372-D16) that form strong electrostatic interactions between ABI1 and ROP11. Notably, K404 in ABI1 directly interact with the phosphate group in GTP of ROP11 (Fig. S11B), which could be the key interaction that promotes the formation of such complex. However, the GTP is largely blocked by the complex, which would not be exposed to GTP/GDP exchange by GEF enzyme. We also identified the association pathway to form the state 240 from our CG MD simulations (Fig. S12). For state 3, the switch I, switch II and insert region are not involved in the ABI1-ROP11 interface. In addition, both N-terminal and C-terminal of ROP11 interact with ABI1, which is unlikely to be physical (Fig. S13). Take altogether, the state 3 is less likely to be the true complex compared to the state 160 and the state 240.

### 2.5 Residue interaction energy and protein energy network analysis reveal the key residue pairs in state 160 and state 240

We further sought to identify the key residue pairs with significant contributions to the interaction energy between ABI1 and ROP11 in both the state 160 and the state 240. We computed the ensemble-average non-bonded interaction energies between the residue pairs in ABI1 and ROP11 from MD trajectories according to the force field parameters used in the MD simulations.^60^ After we obtained the mean interaction energies (MIE), we computed the residue-residue MIE and correlation matrix for both the state 160 (Fig. 6A,B) and the state 240 (Fig. S14A,B). In addition, protein energy network (PEN) was constructed by considering individual residues as nodes and MIEs between residue pairs as the ‘weight’ for the edges that connect these residue nodes. Using PEN, node-based network metrics including degree and betweenness-centralities (BC) were obtained to assess the importance of each residue in terms of protein stability (Fig. 6C and Fig. S14C,D). Specifically, degree measures the number of edges connected to a respective residue and BC measures how frequently this residue occurs in all shortest paths between all other residues. Using the metrics including MIE, correlation, degree and BC, we can identify the residues in ABI1 and ROP11 that are important in forming the complexes in the state 160 and the state 240.

**Figure 6:**
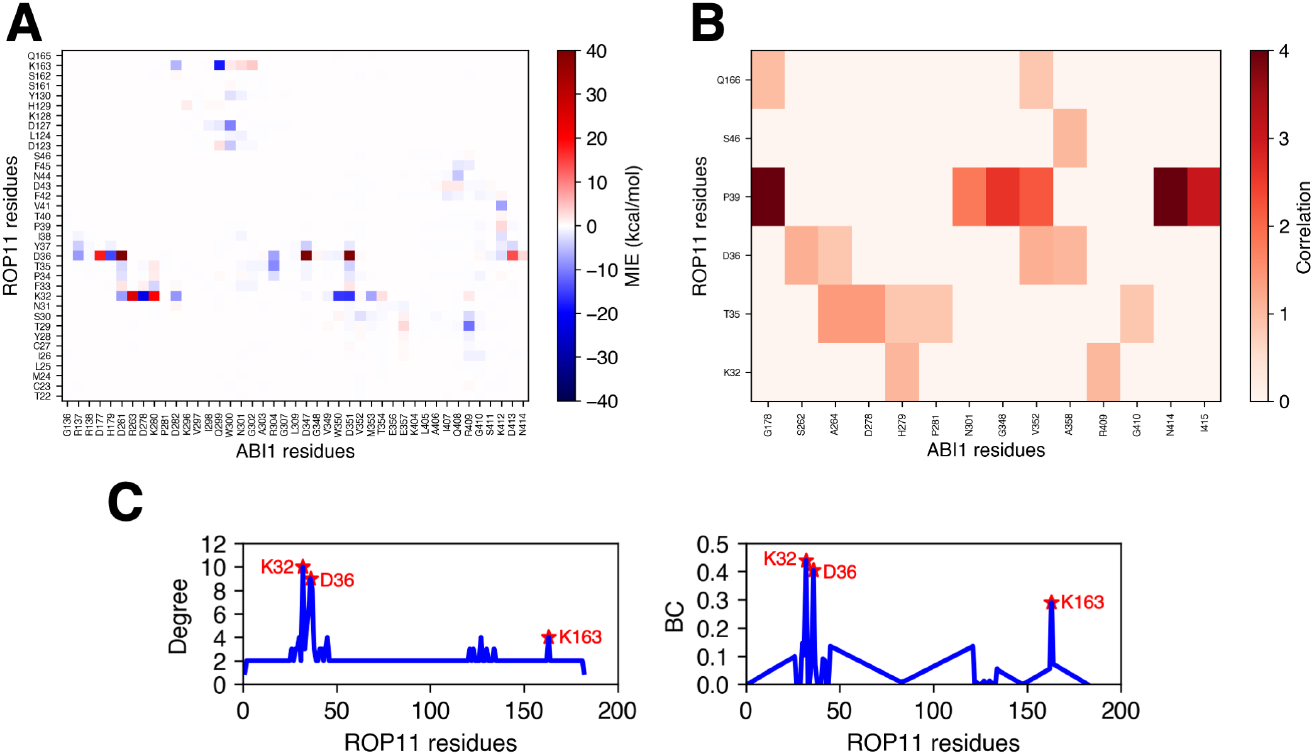
Identification of critical residue-residue pairs in state 160. (A) Mean interaction energy (MIE) matrix and (B) residue correlation matrix for the residue-residue pairs between ABI1 and ROP11. (C) Network analysis reveals the degree and betweenness-centrality (BC) of the residues in ROP11. The residues K32, D36 and K163 in ROP11 have the highest degree and BC, indicating their critical role in stabilizing the state 160.

For the state 160, the MIE matrix (Fig. 6A) highlights the residue pairs with favorable (negative MIE) and unfavorable (positive MIE) contributions to form the complex. The top 20 residue pairs with the lowest MIE and their MIE values are summarized in Table S4. Notably, the three residues in ROP11, including K32, K163 and D36, strongly interact with ABI1 and have the largest contributions to binding energy (Fig. 4E). We note that D36 also interacts with Mg^2+^ at the catalytic site of ABI1, which is expected to have even stronger interaction compared to residue-residue interactions (Fig. 4E). In addition, the three residues and their neighboring residues are correlated with the residues in ABI1 (Fig. 6B), and have relative higher degrees and BCs (Fig. 6C). For the state 240, the charged residue pairs, including R189-D127, E190-K128, K372-D16, K391-D133, and K371-D68, have the largest contributions to form the complex (Table S5, Fig. S14). In addition, K404 in ABI1 interacts with the phosphate groups of GTP in the state 240, which is expected to have favorable interaction energy. Overall, residue interaction energy and protein energy network analysis enables the identification of key residue pairs that are involved in forming the state 160 and the state 240, which can be further validated experimentally.

## 3 Discussion

PPIs play a vital role in plant hormone signal transduction and other biological processes in plants. The majority of PPIs are specific in both their binding targets and the binding sites. It is therefore critical to elucidate the binding partner of a target protein and the molecular details of their interaction, in order to better understand and engineer these biological processes. Despite recent advances in a variety of experimental techniques for determining three-dimensional structures of proteins and complexes, it remains, in many cases, challenging to obtain high-resolution protein complex structures. In particular, high-quality structural information for plant proteins and complexes is scarce.

Molecular modeling techniques are powerful computational tools to complement experiments for structural modeling of PPIs. With the rapid progress in high-performance computing and MD software, one can perform microsecond-to millisecond-long timescale MD simulations to study complex protein dynamics and function at a high spatial-temporal resolution. In contrast to protein-protein docking, MD simulations can capture the dynamic nature of proteins as well as protein interactions with organic ligands and cofactors. However, the application of MD simulations in accurate modeling of PPIs remains to be limited by available computational power and inherent complexity of PPIs due to vast configuration space of protein complexes.

In this study, we have integrated long timescale coarse-grained and all-atom MD simulations for predicting the complex structure of ABI1 and ROP11, which are involved in ABA signaling network. Combining CG MD simulations and Markov state model analysis, we were able to adequately sample the complex configuration space and identify a small number of candidate configurations. The candidate complexes were further refined through all-atom MD simulations and differentiated through binding free energy evaluation. We obtained two candidate complex structures, both the state 160 and the state 240, that require experimental information to validate. By performing the residue interaction energies and protein energy network analysis on all-atom MD trajectories, we identified the critical residues in both ABI1 and ROP11 which contribute favorably to forming the state 160 and the state 240, respectively. From our CG MD simulations, we reported not only the near-native structure of ABI1 and ROP11 but also the association pathway to form the native complex. Overall, this study demonstrates the powerful framework of integrating a range of molecular modeling techniques for predicting protein complex structures.

In conclusion, our study has provided key structural insights into negative regulation of ABA signaling through molecular interactions between protein phosphatase PP2C and small GTPase. The structural information unraveled in this study helps improve molecular understanding of ABA signal transduction mechanism, and can potentially create new avenues to engineer ABA signaling pathway to regulate ABA-mediated responses. From a broad perspective, our computational framework used in this study can be extended to study other PPIs involved in a variety of biological processes. As MD methodology continues to develop, including improved force field accuracy and integration of sequence co-evolution ^61–63^ and experimental information,^45,64^ we expect that molecular simulations can be increasingly useful in understanding and engineering plant proteins and complexes.

## 4 Acknowledgments

The authors acknowledge the support from the Blue Waters sustained-petascale computing project, which is funded by the National Science Foundation (awards OCI-0725070 and ACI-1238993) and the state of Illinois. D.S. acknowledges the support from Foundation for Food and Agriculture Research via the New Innovator Award in Food & Agriculture Research. C.Z. acknowledges the support by 3M Corporate Fellowship and Glenn E. and Barbara R. Ullyot Graduate Fellowship from the University of Illinois at Urbaba-Champaign.

## Supporting Information Available

The Supporting Information is available free of charge at http://pubs.acs.org.

## Data and Software Availability Statement

Simulation trajectories for the coarse-grained and all-atom simulations, as well as data for Replica Exchange Umbrella Sampling for free energy calculations are available at https://uofi.box.com/s/0luwjpu7mbg0mvs1gzaonzijixtm8ww4. Markov state model and clustering objects along with normalized features are available at https://doi.org/10.5061/dryad.h9w0vt4tx.

