## Supporting Information for "Structural Basis for Negative Regulation of ABA Signaling by ROP11 GTPase"

### Methods

#### Homology modeling and structure refinement of ABI1 and active ROP11 GTPase

The initial structures of AtABI1 and the active AtROP11 GTPase were modeled via homology modeling using Swiss-Model.[S1] The crystal structure of AtABI2 (PDB ID: 3UJK[S2]) was used as the template for modeling ABI1 (sequence identity and similarity between ABI1 and ABI2: 87.17% and 57%). Two $Mg^{2+}$ ions bound to the catalytic site of ABI1 were also included in the modeled ABI1. The crystal structure of active OsRAC1 (PDB ID: 4U5X[S3]) was used as the template for modeling ROP11 (sequence identity and similarity between OsRAC1 and ROP11: 82.95% and 56%). The non-hydrolyzable GTP analog GMPPNP and $Mg^{2+}$ in 4U5X were added to the modeled AtROP11 GTPase structure. GMPPNP was subsequently replaced by GTP molecule to obtain the initial homology model of active AtROP11.

Explicit solvent molecular dynamics (MD) simulations were performed to refine the modeled structures of ABI1 and ROP11. The initial structures were solvated using tleap in Ambertools 15.[S4] Extra $Na^{+}$ and $Cl^{-}$ (150 mM) were added to neutralize the system and mimic physiological environment. The parameters for GTP were obtained from AMBER parameter database (http://research.bmh.manchester.ac.uk/bryce/amber/) and Amber ff14SB force field was used for ABI1 and ROP11. The AMBER force field parameters for monovalent and divalent ions available in AMBER 18 software[S4] were used. The system was subjected to 10000 steps of minimization and subsequent 1 ns equilibration at 300 K. Production simulations were launched from the equilibrated configuration and run for 100 ns. The simulations were run in isothermal-isobaric (300 K, 1 atm) ensemble using an integration time step of 2 fs. The temperature was controlled using Langevin dynamics[S5] with the collision frequency of 2 $ps^{-1}$ and the pressure was maintained using Monte Carlo barostat. Periodic boundary condition was applied in all MD simulations. The particle-mesh Ewald method was used to treat the electrostatic interactions, along with a 10 Å cutoff distance for van der Waals interactions.[S6] The SHAKE algorithm[S7] was applied to constrain the length of covalent bonds involving hydrogen atoms.

### Coarse grained (CG) molecular dynamics (MD) simulations of ROP11 and ABI1 association

The atomic structures of ABI1 and ROP11 obtained after MD refinement were used to construct CG structural models and run CG MD simulations using Martini force field.[S8,S9] For ROP11, Q66L mutation was performed using Pymol to mimic a constitutively active ROP11 (CA-ROP11).[S10,S11] In such way, GTP and $Mg^{2+}$ were not included in CA-ROP11 structure. The script Martinize.py (version 2.6, http://cgmartini.nl/index.php/tools2/proteins-and-bilayers) was used to convert the atomic structures of CA-ROP11 and ABI1 to CG bead models and also generate the Martini topology files in Gromacs format. In short, CG bead models were produced according to 4 heavy atoms to 1 bead mapping and different types of beads were defined to represent different subgroup of atoms. Elastic network was applied in combination with Martini model to conserve secondary, tertiary and quartenary structures without sacrificing realistic dynamics of the proteins.[S12] *ElNeDyn* network was used to generate the topology files of the proteins.[S12]

Next, the CG structural models of ABI1 and CA-ROP11 were randomly placed far away from each other to generate 50 configurations of ABI1/CA-ROP11 complexes, with their center-of-mass distance at least 25 Å. The 50 complex structures were solvated using an equilibrated CG water box via Gromacs solvate command. Also, 30 CG $Na^{+}$ and 31 $Cl^{-}$ beads were added to neutralize the system. The system was subjected to 0.2 ns equilibration in Gromacs.[S13] Production simulations were launched from the equilibrated configurations. Adaptive sampling approach was used to efficiently sampling the association process between ABI1 and CA-ROP11.[S14] In the initial round of simulations, the simulations started from 50 configurations. In the next rounds of simulations, the previously sampled complex conformations were clustered into a certain number of states. A portion of the states with the lowest populations were selected to start new simulations, with the initial velocities randomly assigned according to Boltzmann distribution.

The CG MD simulations were performed with Gromacs 2019[S13] in isothermal-isobaric (300 K, 1 atm) ensemble using an integration time step of 20 fs. The temperature control was achieved with the velocity rescale (V-rescale) thermostat using a coupling constant of the order of 1 ps. The pressure control was realized with the Parrinello-Rahman barostat, with a coupling constant of 16 ps and compressibility of $3*10^{-4}$ $bar^{-1}$. Periodic boundary conditions were applied in all MD simulations. The particle-mesh Ewald method was used to treat the electrostatic interactions, along with a 11 Å cutoff distance for van der Waals interactions.[S6] In total, 5 rounds of adaptive sampling simulations were performed, leading to an aggregate of 10.05 ms simulations (Table S2).

### Markov state model analysis of simulation data on CA-ROP11 and ABI1 association

Markov state model (MSM) was used to analyze the conformational ensemble of ABI1 and CA-ROP11 resulted from CG MD simulations. MSM is a powerful method to analyze large-scale MD simulation data on biomolecules.[S15,S16] Generally, MSMs characterize biomolecule conformation ensemble as discrete conformational states and inter-state transition probabilities between them.[S15,S16] The equilibrium probabilities of individual states can then be estimated from MSM, allowing us to gain insights into the probable configurations of ABI1/CA-ROP11 complex.

The discretization of conformational space was achieved by identifying a set of structural features to represent individual configurations and then clustering these configurations into a certain number of states. In total, 6 structural metrics were calculated to differentiate various configurations, including the center-of-mass distance between ABI1 and CA-ROP11 as well as 5 angles or dihedral angles that describe the relative orientation and position of ABI1 and CA-ROP11 (Fig. S2). Time-lagged independent component analysis was performed to identify several slowest degrees of freedom (tICs) resulted from linear combination of the original metrics.[S17] K-means clustering were then performed on the identified slowest motions to cluster the CG configurations into a certain number of states. MSMs were then constructed based on the clustering results. The matrix of transition probabilities was determined with a lag time $\tau$ using maximum likelihood approximation. The optimal lag time ($\tau$) was chosen based on the convergence of the implied timescales of MSMs and the number of clusters ($N$) as well as the number of tICs ($M$) were optimized via cross validation ranked by the variational GMRQ objective function (Fig. S3).[S18] All MSMs were constructed using the MSMBuilder 3.4[S19] and the MSM hyperparameters were optimized using the Osprey software.[S20] The final MSM hyperparameters were 600 clusters, 4 tICs, and 800 ns lag time.

### All atom MD simulations for structural refinement of 23 candidate complex structures

The 25 MSM states with the largest equilibrium populations were converted back to atomic structural models using the script backward.py (http://www.cgmartini.nl/index.php/downloads/tools/240-backward).[S21] The backward mapping was performed using the wrapper initram.sh, which calls backward.py and subsequently relaxes the resulting structure through energy minimization and molecular dynamics based relaxation. Due to the possible inaccuracy of structural models from backward mapping, we aligned the atomic structures of ABI1 and GTP-bound ROP11 to these 25 complexes and sought to use MD simulations to further refine these candidate complex structures. We observed that 3 states among the top 25 states were similar so that we merged the 3 states, resulting in 23 states to be optimized.

The atomic structures of the 23 candidate complexes were solvated with water molecules and counterions. Two parallel MD simulations were launched from the equilibrated configurations with Amber 18[S4] and each was run for 240 ns to relax the atomic structures. The same set of force field parameters and simulation protocols as in the step of structural refinement of homology models were used to run these simulations. After the simulations were finished, the root mean square deviation of these complex structures and the center-of-mass distance between ABI1 and ROP11 were calculated to characterize the relative stability of the 23 complex structures. Based on these simulation data, we further excluded 10 complex structures, leading to 13 remaining candidate complex structures.

### Replica exchange umbrella sampling MD simulations for potential of mean force calculations

To further differentiate the 13 candidate complex structures, we sought to calculate potential of mean force (PMF) for separating ABI1 and ROP11 in these different complex structures. The PMFs were determined by separating two proteins in the presence of a series of conformational, positional and orientational restraints and then performing replica exchange umbrella sampling (REUS) MD simulations to estimate the conformational probability distribution with respect to separation distance $r$. By applying these restraints, it effectively reduce the configuration entropy of complex and accelerate the convergence of separation PMF. In total, there were 7 harmonic restraints applied during separation, including two relating to conformational changes in ABI1 and ROP11 (denoted by subscripts $\mathrm{B}_{ABI1}$, $\mathrm{B}_{ROP11}$) and 5 relating to relative position and orientation between two proteins. The conformational restraints acting on heavy-atom RMSD of the backbone in each protein ($\mathrm{B}_{ABI1}$, $\mathrm{B}_{ROP11}$) restrict protein structural fluctuations and deviations from the starting associate state. Three angular restraints acting on the relative orientation of two proteins (denoted by subscript $o$, including $\Theta$, $\Phi$, $\Psi$) and two on the relative position (denoted by $a$, including $\phi$, $\theta$) limit the configuration space when two proteins are separated.

REUS MD simulations[S22] were used to calculate the separation PMFs of the 13 candidate complex structures. REUS simulations were performed using NAMD 2.13 software.[S23] The complexes were solvated with a sufficiently large water box which encompasses both ABI1 and ROP11 when fully separated. After 20000 steps energy minimization, each system was equilibrated in NPT ensemble for 1 ns. An additional 200 ps simulation for each system was performed to measure the average values of all restrained CVs in the associated state. Next, steered MD simulations were performed to slowly increase the center-of-mass distance between ABI1 and ROP11 ($r$) by 15 Å over 8 ns, in order to generate starting structures for REUS MD simulations. From the trajectory, 31 structures with $r$ evenly spaced between the minimum and the maximum distances were chosen as the starting structures for REUS MD simulations (31 windows, 0.5 Å/window). The starting structure in each umbrella window was equilibrated for 1 ns and then each replica was run for 8 ns. The force constants for the restrained CVs were given in Table S3. The separation PMF was estimated using the multistate Bennett acceptance ratio (MBAR) method with pymbar python package.[S24] The trajectories from simulations were sorted and subsampled to ensure uncorrelated samples for estimating PMF. According the separation PMFs, we identified 3 candidate complex structures which show relative higher stability as compared to other states.

### Replica exchange umbrella sampling MD simulations for standard binding free energy calculations

We sought to determine the absolute standard binding free energy ($\Delta G^o_{bind}$) of the 3 candidate complex structures. The PMF-based approach to determining $\Delta G^o_{bind}$ was developed by prior studies.[S25–S27] The detailed theory of this method was thoroughly described in our previous work.[S28] Integration of the separation PMF contributes to the major component of $\Delta G^o_{bind}$. Next, the contributions of adding the conformational and angular restraints on the associated state and removing them from the fully dissociated state to $\Delta G^o_{bind}$ were also computed and then subtracted from the free energy component resulting from separation PMF. Given the equilibrium association constant, $\Delta G^o_{bind}$ is given by

$$\Delta G^o_{bind} = -\beta^{-1} ln(K_{eq} C^o) \tag{1}$$

where $\beta$ is the reciprocal of the product of gas constant and temperature, and $C^o$ is standard concentration of 1 M, which is 1/1661 Å$^3$. $K_{eq}$ can be expressed as in equation 2.

$$\begin{aligned} K_{eq} = S^* I^* e^{-\beta[(G^{bulk}_{B_{1,c}} - G^{site}_{B_{1,c}}) + (G^{bulk}_{B_{2,c}} - G^{site}_{B_{2,c}})]} \\ * e^{-\beta[(G^{bulk}_{o} - G^{site}_{o}) - G^{site}_{a}]} \end{aligned} \tag{2}$$

The term $S^*$ addresses the removal of positional restraints ($\theta$, $\phi$) on one protein, which is separated from the other protein along $r$ instead of free diffusion. $S^*$ is given by

$$S^* = r^{*2} \int_0^{\pi} d\theta sin\theta \int_0^{2\pi} d\phi e^{-\beta u_a(\theta,\phi)} \tag{3}$$

where $r^*$ is a point far from the binding site and $u_a = u_\theta + u_\phi$. The term $I^*$ is given by

$$I^* = \int_{site} dr e^{-\beta[W(r) - W(r^*)]} \tag{4}$$

which includes the contribution from the separation PMF $W(r)$.

The exponential terms in equation 2 are related to adding the restraints on the associated state (denoted by site) and removing the restraints from the dissociated state (denoted by bulk). These terms can be determined by calculating a series of PMFs for individual restraints and integrating over the PMFs. REUS MD simulations along the restrained CVs were performed to calculate the free energy contributions of adding those restraints to the associated state and removing them from the dissociated state.

To generate a wide range of starting structures for each restrained CV, we performed 6 ns temperature-accelerated MD simulations, coupling the respective CV to a dummy particle experiencing a temperature of 2500 K. The starting structures for REUS windows were chosen from the accelerated MD trajectory, with a window size of 0.05 Å for conformational restraints and 1° for angular restraints. Each replica was equilibrated for 1 ns and then run for 8 ns. The force constants used were 1000 kcal·mol$^{-1}$·Å$^{-2}$ for RMSD restraints and 2.5 kcal·mol$^{-1}$·deg$^{-2}$ for angular restraints.

The PMFs were determined using MBAR. For some of the RMSD restraints, we performed additional targeted MD simulations and umbrella sampling simulations to expand the ranges of PMFs and ensure the convergence of PMFs. Given the PMFs, the contributions of adding and removing restraints to $\Delta G^{o}_{bind}$ can be calculated via numerical computations. Please refer to our previous work for more details.[S28]

### Residue-residue Interaction energies and protein energy network analysis

To identify the critical residue pairs for intermolecular interaction between ABI1 and ROP11, we performed residue-residue interaction energies and protein energy network analysis on MD trajectories using gRINN version 1.1.0.hf1 (https://grinn.readthedocs.io/en/latest/).[S29] gRINN computes pairwise residue non-bonded interaction energies by repeatedly calling the MD simulation engine in GROMACS 2019.[S13] Default values of 60% percent cutoff and 20 Å filtering distance cutoff were used, which implied that only the residue pairs between ABI1 and ROP11 whose center-of-mass distances are less than 20 Å in at least 60% of trajectory frames will be included in the calculation. In total, 250 frames of all-atom MD trajectories for the state 160 and the state 240 were used for the calculations. Residue pairwise correlations and protein energy network (PEN) were also computed using gRINN. Protein energy network (PEN) was constructed by considering individual residues as nodes and mean interaction energies between residue pairs as the 'weight' for the edges that connect these residue nodes.

Using PEN, node-based network metrics including degree and betweenness-centralities were obtained to assess the importance of each residue in terms of protein stability.

Table S1: Overview of computational simulations and analysis methods performed in this study.

| Step | System | Method | Force Field | Software | Size (Å$^3$) | Ensemble | Sim. Time |
|---|---|---|---|---|---|---|---|
| 1 | ROP11 active structure | Homology modeling | - | Swiss-Model | - | - | - |
| 2 | ROP11 structure refinement | All-atom MD | Amber ff14SB | Amber 18 | 67*67*81 | NPT | 100 ns |
| 3 | CA-ROP11 (Q66L)-ABI1 association | Adaptive sampling, CG MD | Martini | Gromacs 2019 | ∼140*140*140 | NPT | 10.05 ms |
| 4 | Analysis of complex ensemble | Markov state model | - | MdTraj, MSMBuilder | - | - | - |
| 5 | Structural refinement of top 23 candidates | All-atom MD | Amber ff14SB | Amber 18 | ∼120*120*120 | NPT | 240*23*2 ns |
| 6 | PMF and binding free energy calculations | REUS MD | Amber ff14SB | NAMD 2.13 | ∼80*95*138 | NPT | >=8 ns/window |
| 7 | Residue interaction energies and protein energy network analysis | - | Amber ff14SB | gRINN | - | - | - |

Table S2: Summary of the adaptive coarse grained MD simulations of the association process between ABI1 and CA-ROP11 (Q66L). In the initial round of simulations, the simulations started from 50 configurations where ROP11 and ABI1 were randomly placed far away from each other. In the next rounds of simulations, the previously sampled complex conformations were clustered into a certain number of states. A portion of the states with the lowest populations were selected to start new simulations, with the initial velocities randomly assigned according to Boltzmann distribution.

| Round | Parallel simulations | Simulation time ($\mu$s) | Aggregate (ms) |
|---|---|---|---|
| 1 | 50 | 21 | 1.05 |
| 2 | 100 | 10 | 1 |
| 3 | 200 | 10 | 2 |
| 4 | 200 | 10 | 2 |
| 5 | 400 | 10 | 4 |
| Total simulation time: ~10.05 milliseconds | | | |

Table S3: The restrained CVs and the corresponding force constants used in REUS MD simulations on the separation of candidate ROP11-ABI1 complex structures.

| CV | $k_{force}$ |
|---|---|
| $B_{1,c}$ | 25 kcal·mol$^{-1}$·Å$^{-2}$ |
| $B_{2,c}$ | 25 kcal·mol$^{-1}$·Å$^{-2}$ |
| $\Theta$ | 0.1 kcal·mol$^{-1}$·$^{\circ-2}$ |
| $\Phi$ | 0.1 kcal·mol$^{-1}$·$^{\circ-2}$ |
| $\psi$ | 0.1 kcal·mol$^{-1}$·$^{\circ-2}$ |
| $\phi$ | 0.1 kcal·mol$^{-1}$·$^{\circ-2}$ |
| $\theta$ | 0.1 kcal·mol$^{-1}$·$^{\circ-2}$ |
| $r$ | 25 kcal·mol$^{-1}$·Å$^{-2}$ |

Table S4: The 20 residue-residue pairs between ABI1 and ROP11 with the highest mean interaction energy (MIE) in state 160. MIE was calculated from a 500-ns all-atom MD trajectory using gRINN.[S29]

| ROP11 residues | ABI1 residues | MIE (kcal/mol) |
|---|---|---|
| K32 | D278 | -22.49 |
| K163 | Q299 | -18.18 |
| K32 | D351 | -16.15 |
| K32 | W350 | -15.79 |
| D36 | H179 | -14.58 |
| T29 | R409 | -11.13 |
| D127 | W300 | -9.82 |
| T35 | R304 | -9.20 |
| K32 | D282 | -8.19 |
| D36 | R137 | -8.00 |
| K32 | D261 | -7.74 |
| V41 | K412 | -7.39 |
| D36 | R304 | -7.19 |
| K32 | M353 | -7.15 |
| K163 | D282 | -5.92 |
| N44 | Q408 | -4.40 |
| D123 | W300 | -4.33 |
| Y37 | D347 | -3.99 |
| Y37 | R137 | -3.91 |
| T35 | D351 | -3.57 |

Table S5: The 20 residue-residue pairs between ABI1 and ROP11 with the highest mean interaction energy (MIE) in state 240. MIE was calculated from a 250-ns all-atom MD trajectory using gRINN.[S29]

| ROP11 residues | ABI1 residues | MIE (kcal/mol) |
|---|---|---|
| D127 | R189 | -65.05 |
| K128 | E190 | -32.65 |
| D16 | K372 | -28.93 |
| D133 | K391 | -20.86 |
| D68 | K371 | -13.16 |
| H129 | E190 | -12.78 |
| H134 | K391 | -11.79 |
| V90 | K391 | -6.47 |
| D133 | S146 | -5.66 |
| D68 | K372 | -5.25 |
| G65 | K372 | -4.27 |
| P135 | K391 | -3.92 |
| Y130 | K391 | -2.88 |
| S88 | E400 | -2.72 |
| K128 | N186 | -2.50 |
| Y37 | Q408 | -2.43 |
| D133 | Y129 | -2.4 |
| D127 | A144 | -2.11 |
| D133 | A144 | -2.05 |
| L124 | R189 | -1.72 |

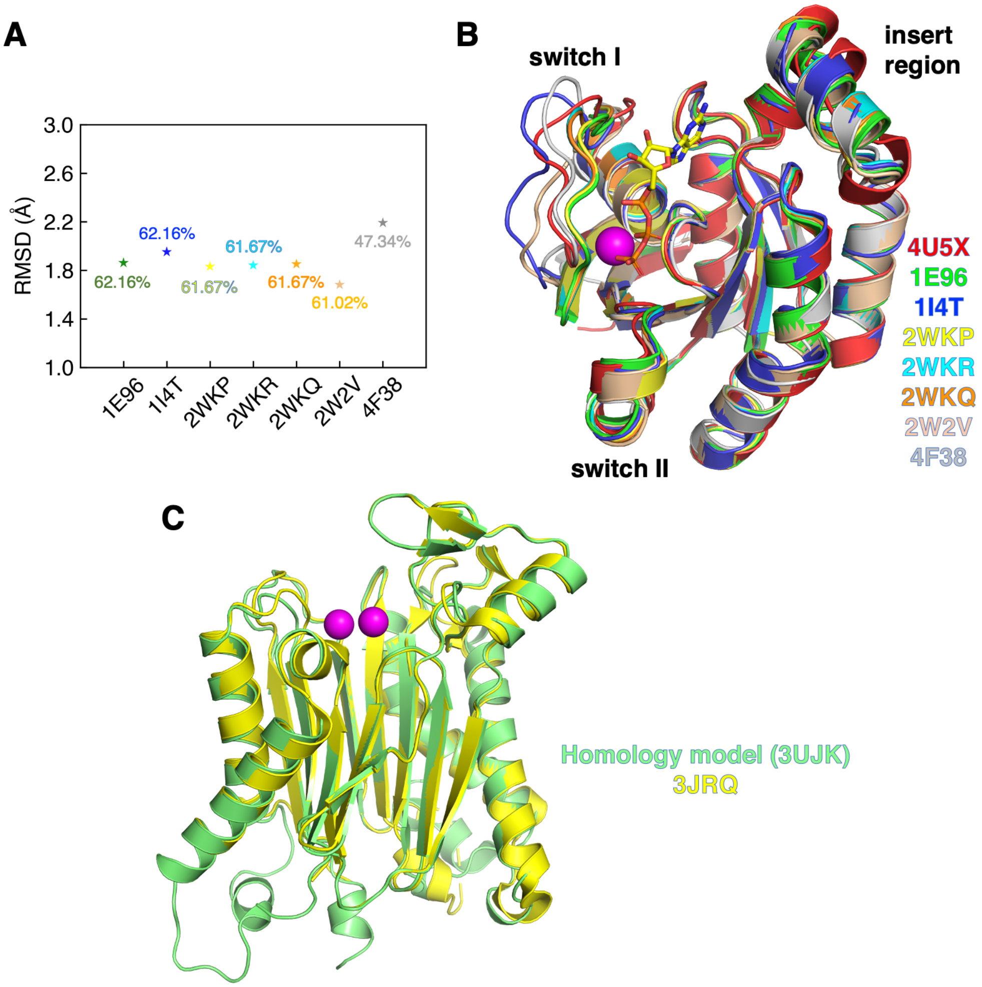


Figure S1: Evaluation of the quality of homology models of AtROP11 and AtABI1. (A) Root mean square deviation between the crystal structures of OsRAC1 (the GTPase template for ROP11, PDB ID: 4U5X) and seven active GTPases with varying degrees of sequence identify to OsRAC1 (PDB IDs: 1E96, 1I4T, 2WKP, 2WKR, 2WKQ, 2W2V, and 4F38) are generally smaller than 2.2 Å. (B) The crystal structures of OsRAC1 and seven GTPases share the same fold, and the switch II and the insert region are highly conserved while the switch I region demonstrates slight variation. As the sequence identity between AtROP11 and OsRAC1 is 82.95%, the homology model of ROP11 using OsRAC1 as the template is of good quality. (C) Structural comparison between the homology model of ABI1 (template PDB ID: 3UJK, lime) and the crystal structure of ABI1 (PDB ID: 3JRQ, yellow). The RMSD between two structures is 1.42 Å.

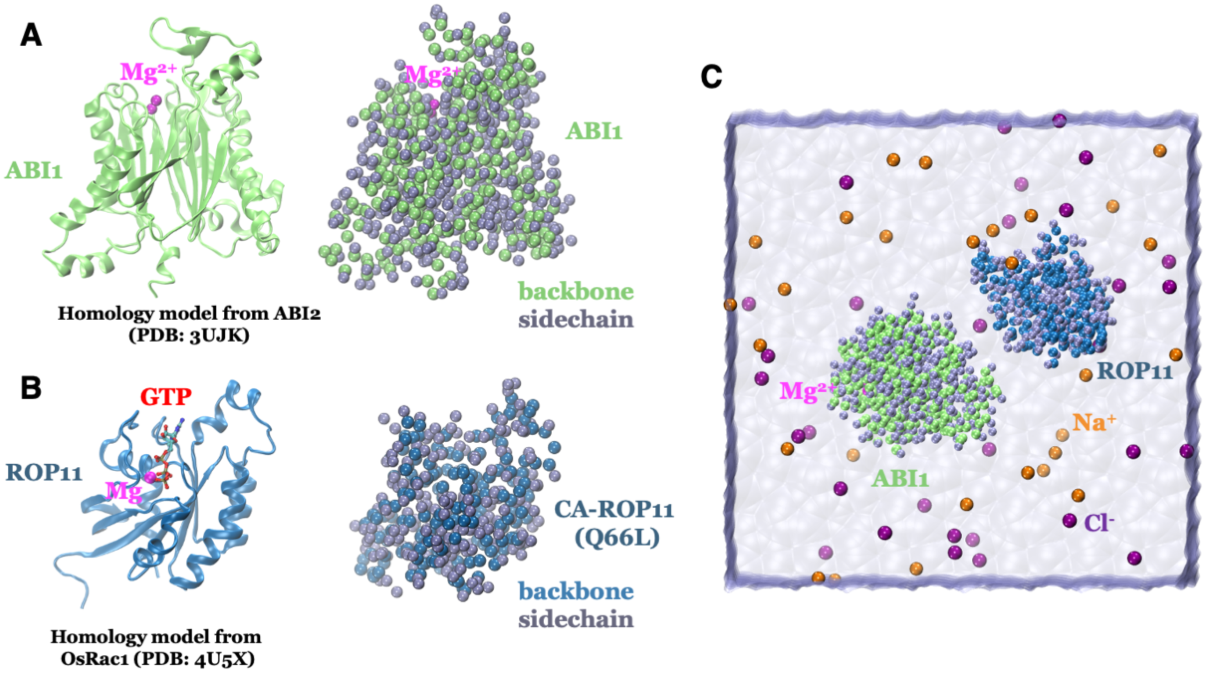


Figure S2: Schematic of CG models of ABI1 and CA-ROP11 and CG MD simulations. Atomic (left) and CG bead (right) representations of (A) ABI1 and (B) CA-ROP11. Backbone beads of ABI1 and CA-ROP11 are shown in lime and blue, respectively. Sidechain beads are shown in purple. (C) A representative view of explicit solvent and periodic CG MD simulation box.

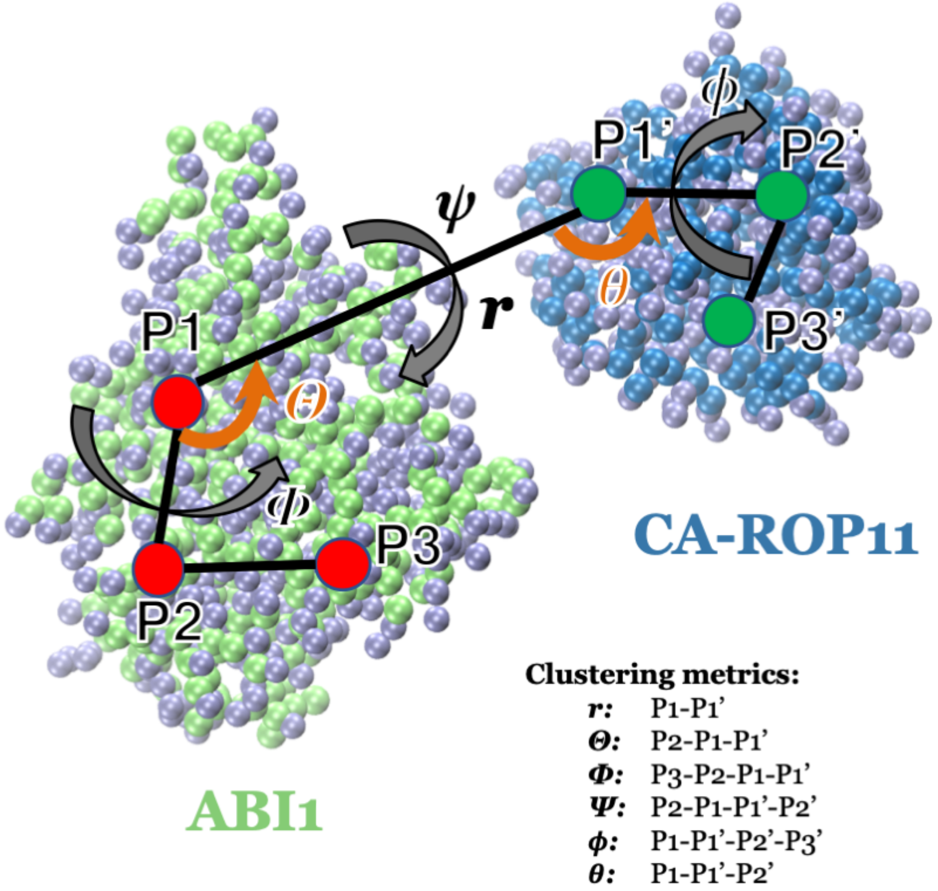


Figure S3: Schematic of structural features used for clustering ABI1/CA-ROP11 complex conformations. In both ABI1 and ROP11, three points (P1, P2 and P3 in ABI1, P1$'$, P2$'$ and P3$'$ in CA-ROP11) were defined as the centers of mass of groups of atoms. The distance between P1 and P1$'$ characterizes the distance between ABI1 and CA-ROP11. The angles $\Theta$ (P2-P1-P1$'$) and $\theta$ (P1-P1$'$-P2$'$) as well as the dihedral angles $\Phi$ (P3-P2-P1-P1$'$), $\phi$ (P1-P1$'$-P2$'$-P3$'$) and $\psi$ (P2-P1-P1$'$-P2$'$) together characterizes the relative position and orientation between ABI1 and CA-ROP11.

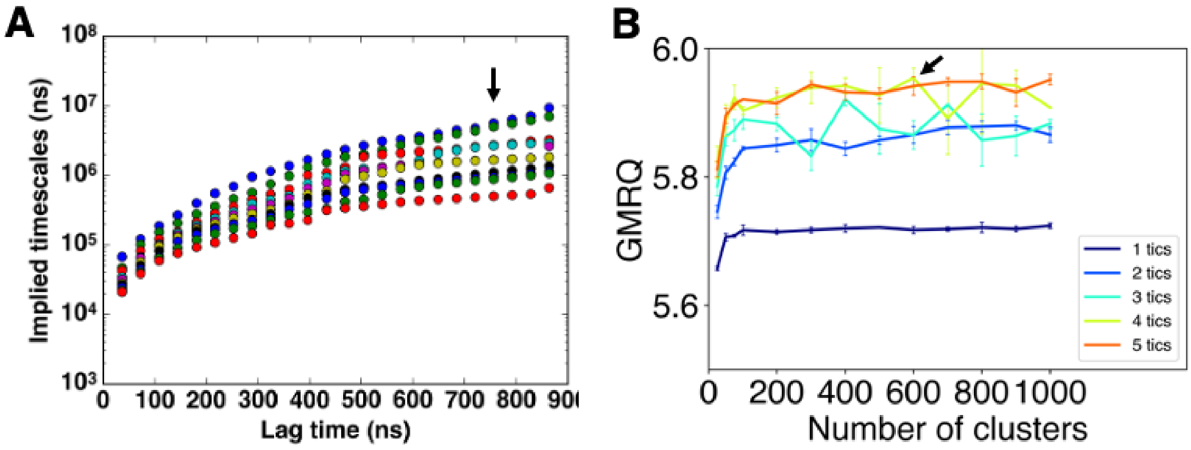


Figure S4: Hyperparameter selection for construction of Markov state models of ABI1/CA-ROP11 association. (A) Convergence of the implied timescales of the slowest processes captured by MSMs with respect to increasing lag times. A lag time of 800 ns was used for the final MSM. (B) GMRQ scores for ranking how well MSMs with various numbers of clusters and tICs capture the slowest processes of ABI1 and ROP11 association. Based on the GMRQ scores, the set of parameters (600 clusters and 4 tICs) was chosen to construct the final MSM.

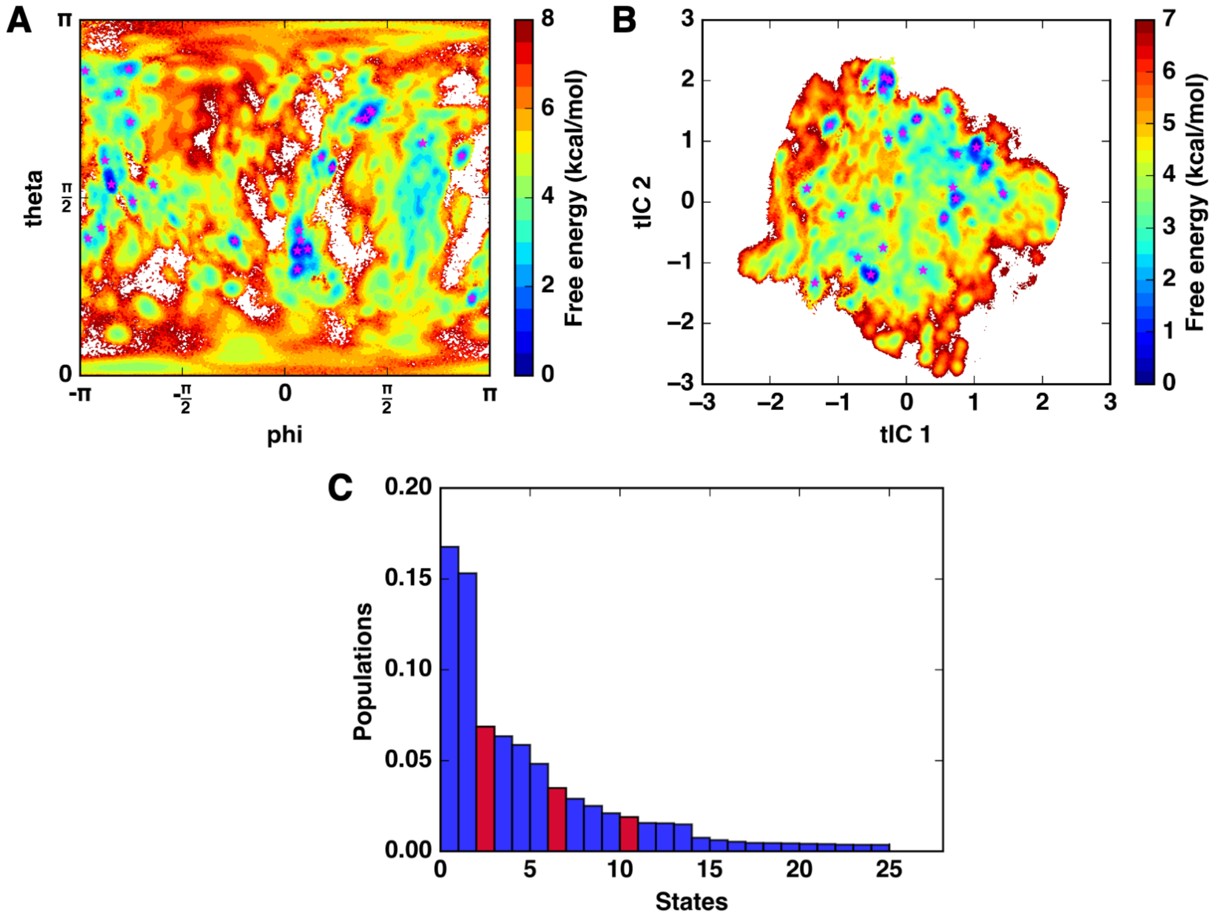


Figure S5: ABI1-ROP11 conformational ensemble sampled from CG MD simulations. Conformational free energy landscape projected onto (A) $\phi$ and $\theta$ and (B) the two slowest degrees of freedom (tIC1, tIC2) resulted from time-lagged independent component analysis of the clustering metrics. Top 25 most populated MSM states are mapped onto the free energy landscape according to their tIC1 and tIC2 values. (C) The equilibrium populations of top 25 states ranked by descending order. Sum of the populations of the top 25 states is greater than 80% of the total populations. The three states with similar complex structures are highlighted in red.

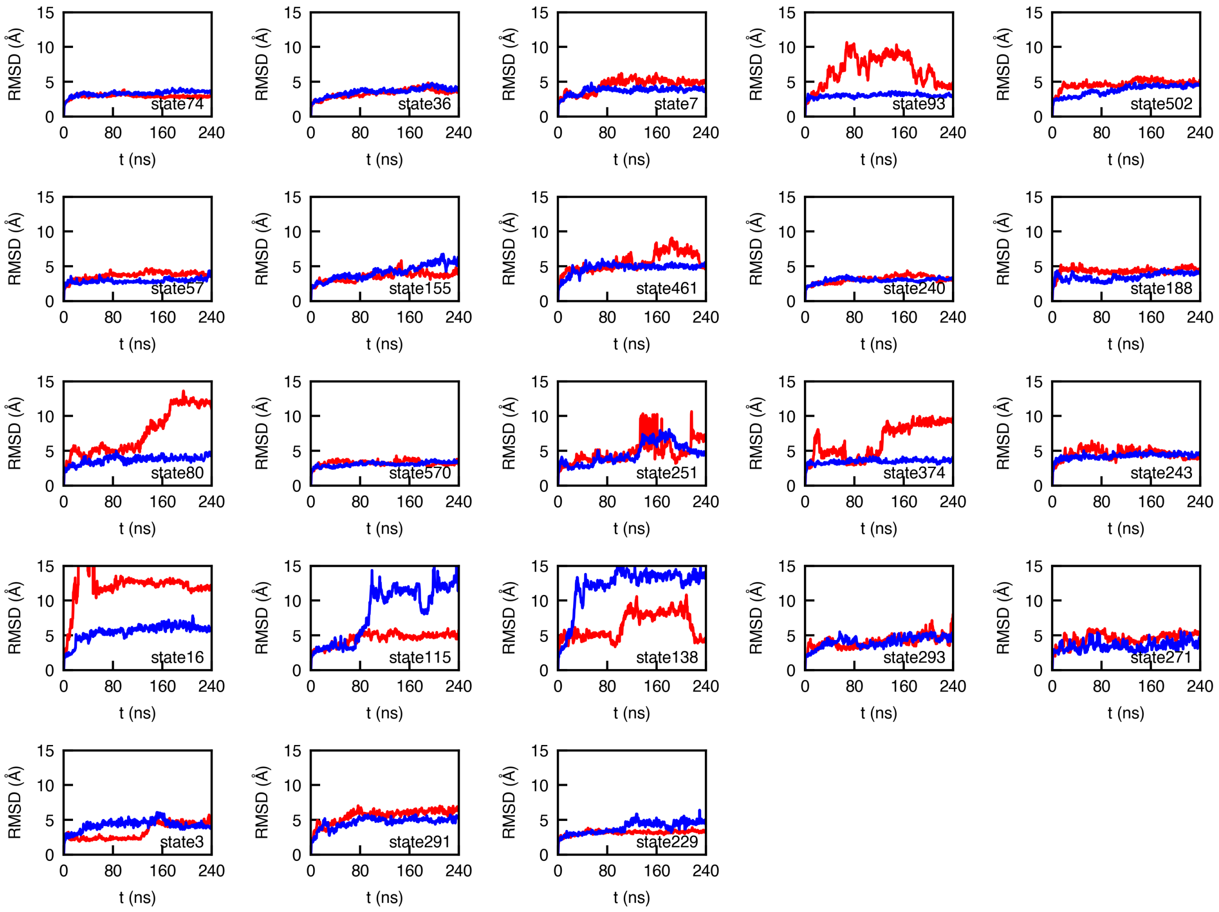


Figure S6: Plots of RMSD, with respect to the first frame of each trajectory, for the 23 candidate complex structures during 240 ns all atom MD simulations.

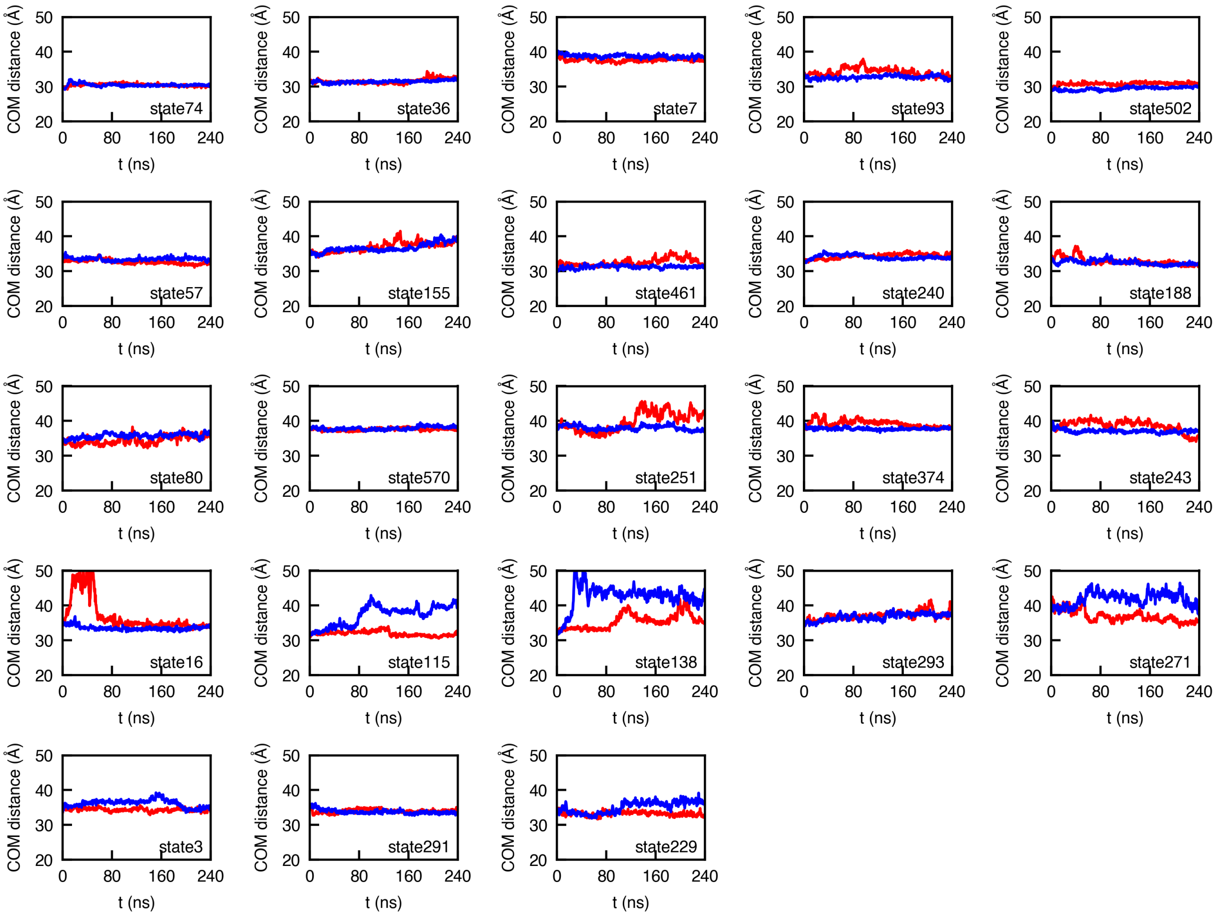


Figure S7: Plots of center-of-mass distance for the 23 candidate complex structures during 240 ns all atom MD simulations.

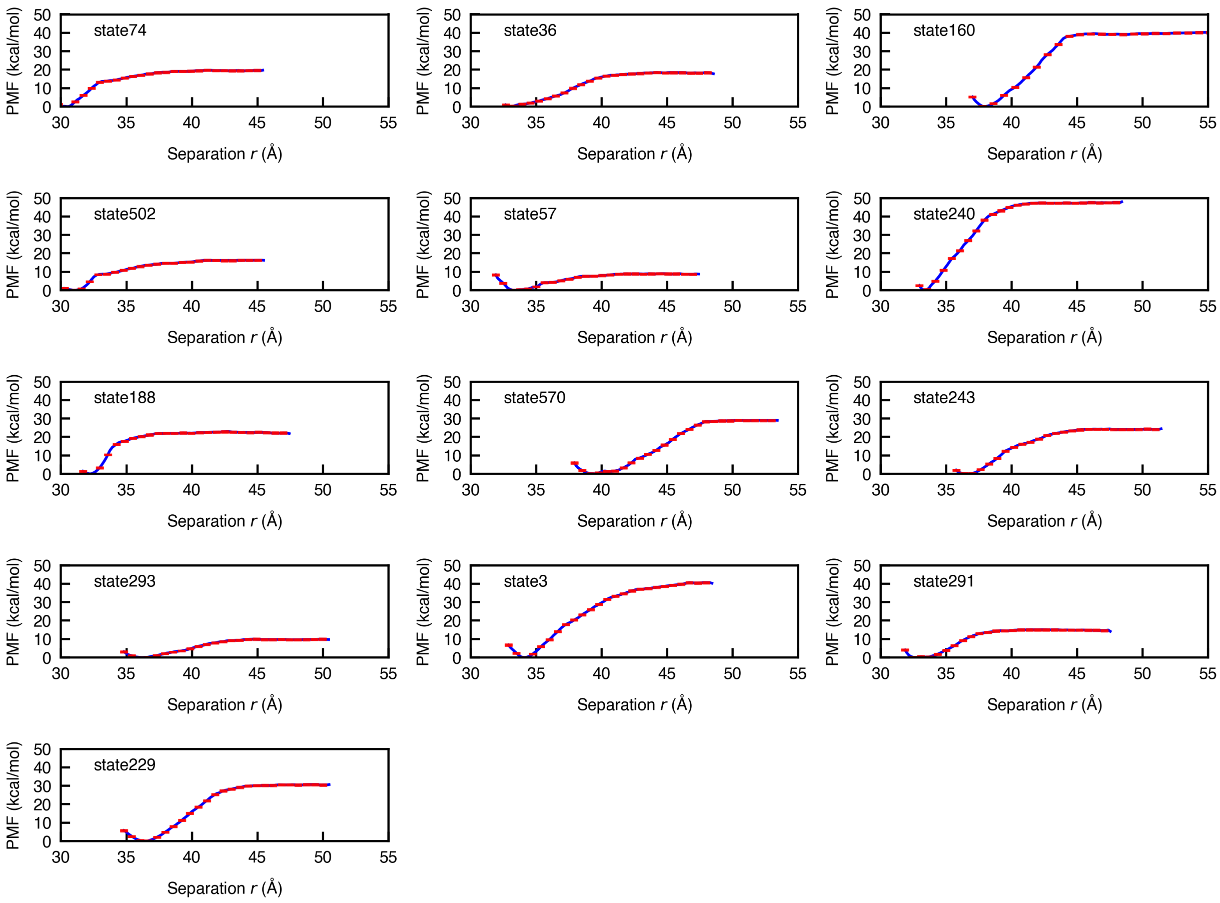


Figure S8: Potential of mean force profiles for separating the top 13 candidate complex structures in the presence of conformational and angular restraints. The error bars on the PMFs are shown. The PMF depths indicate relative stability of the 13 candidate complex structures.

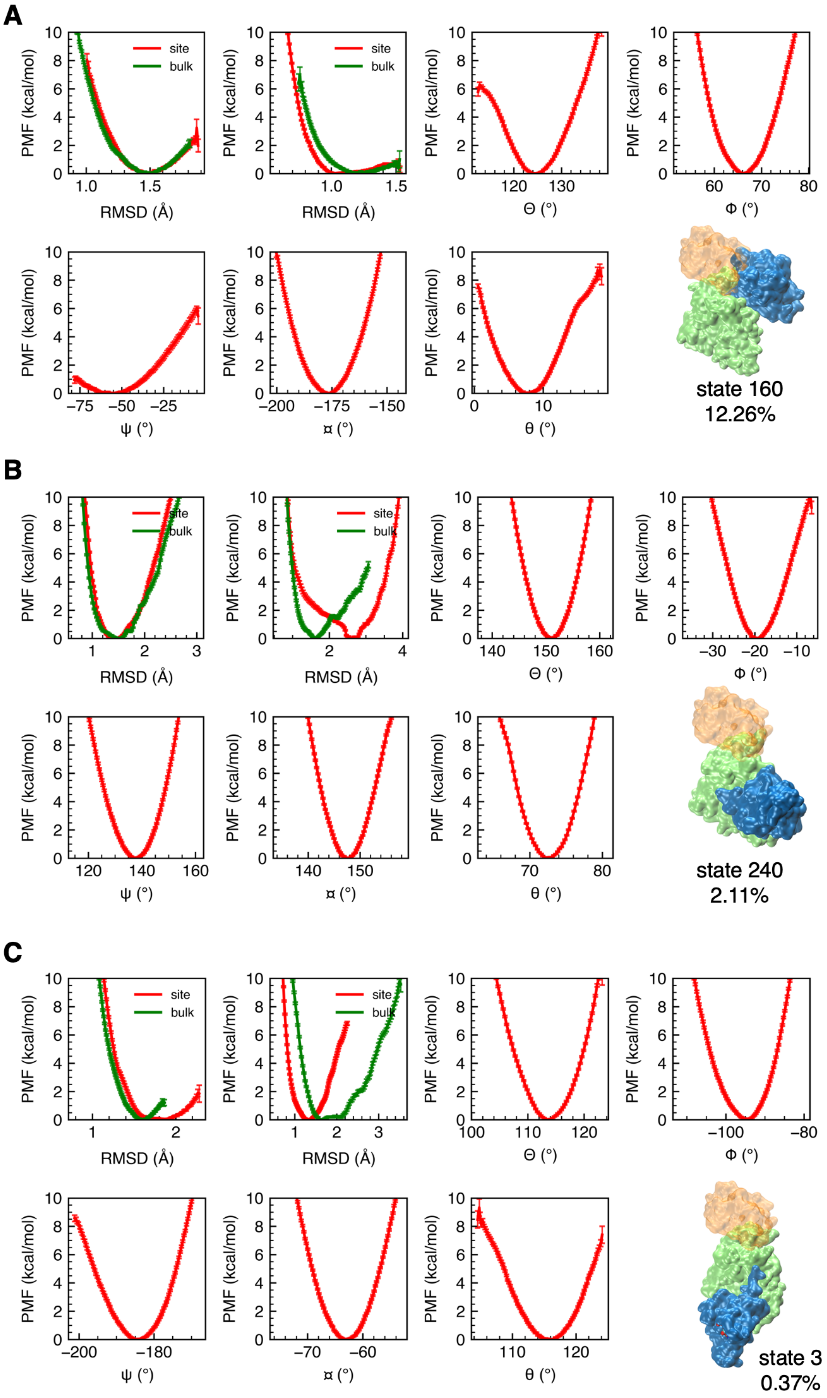


Figure S9: Individual PMFs for all restraint components on (A) state 160, (B) state 240, and (C) state 3. Plots for the conformational and angular restraints $B^{site}_{ABI1,c}$ and $B^{bulk}_{ABI1,c}$, $B^{site}_{ROP11,c}$ and $B^{bulk}_{ROP11,c}$, $\Theta$, $\Phi$, $\psi$, $\phi$ and $\theta$ are shown along with the error bars on the PMFs.

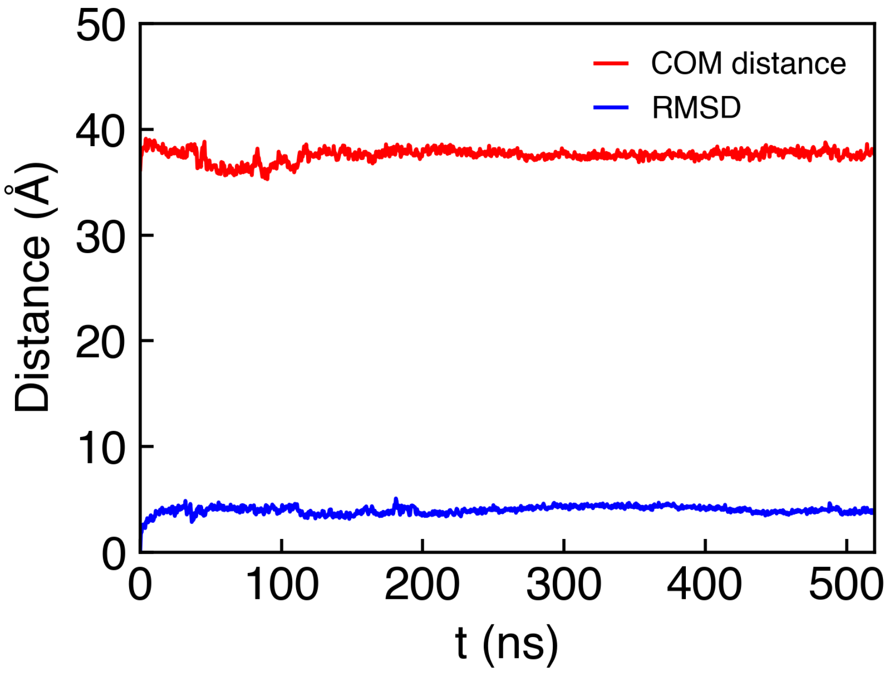


Figure S10: The predicted complex structure (state 160) remains highly stable within 500 ns simulations. RMSD from the initial frame and the center-of-mass distance between ABI1 and ROP11 are plotted with respect to simulation time.

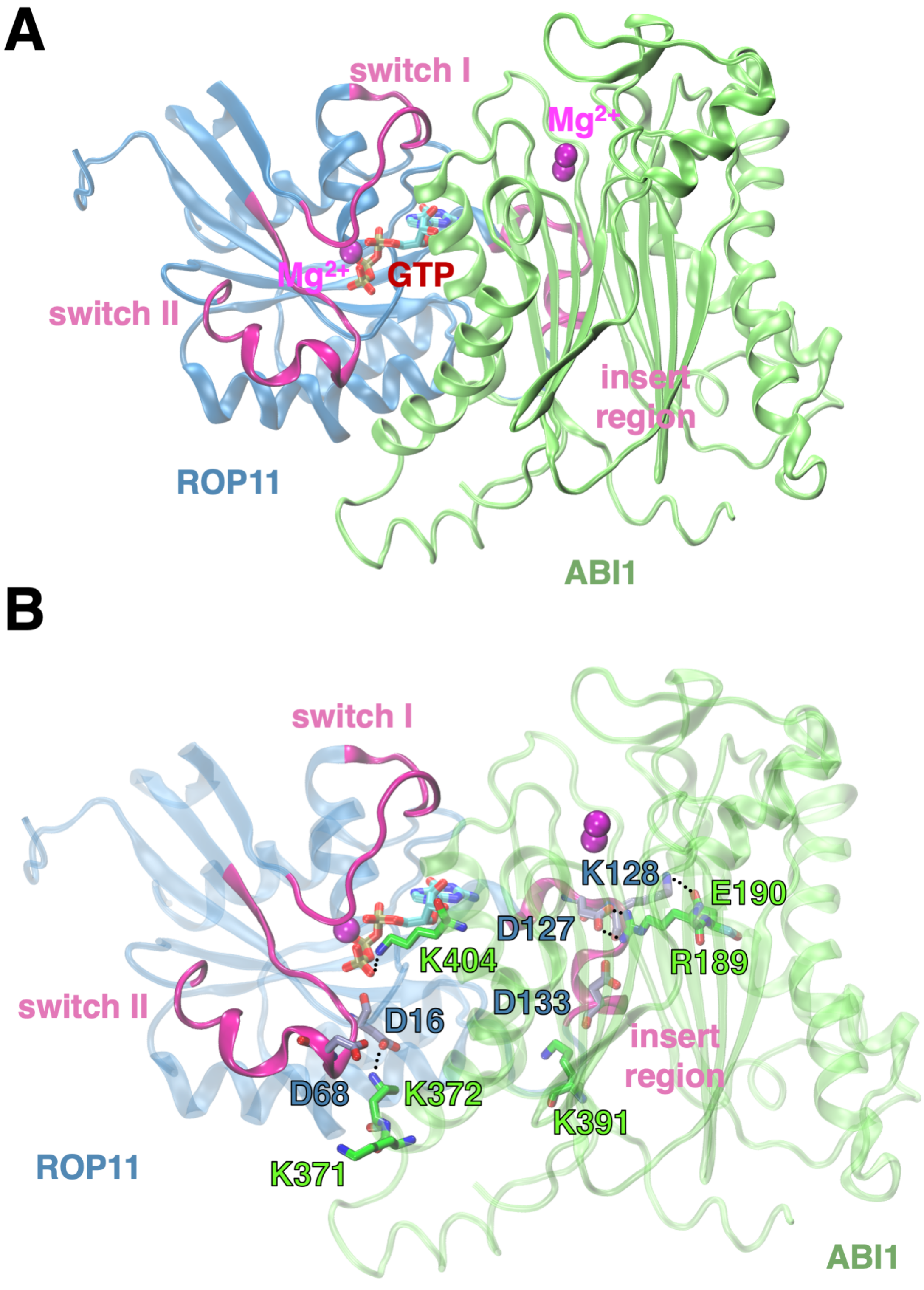


Figure S11: (A) The candidate complex structure of the state 240 and (B) the molecular interactions at ABI1-ROP11 interface in the state 240. The state 240 is stabilized via the electrostatic interaction between both the charged residue pairs in ABI1 and ROP11 and GTP and K404 in ABI1.

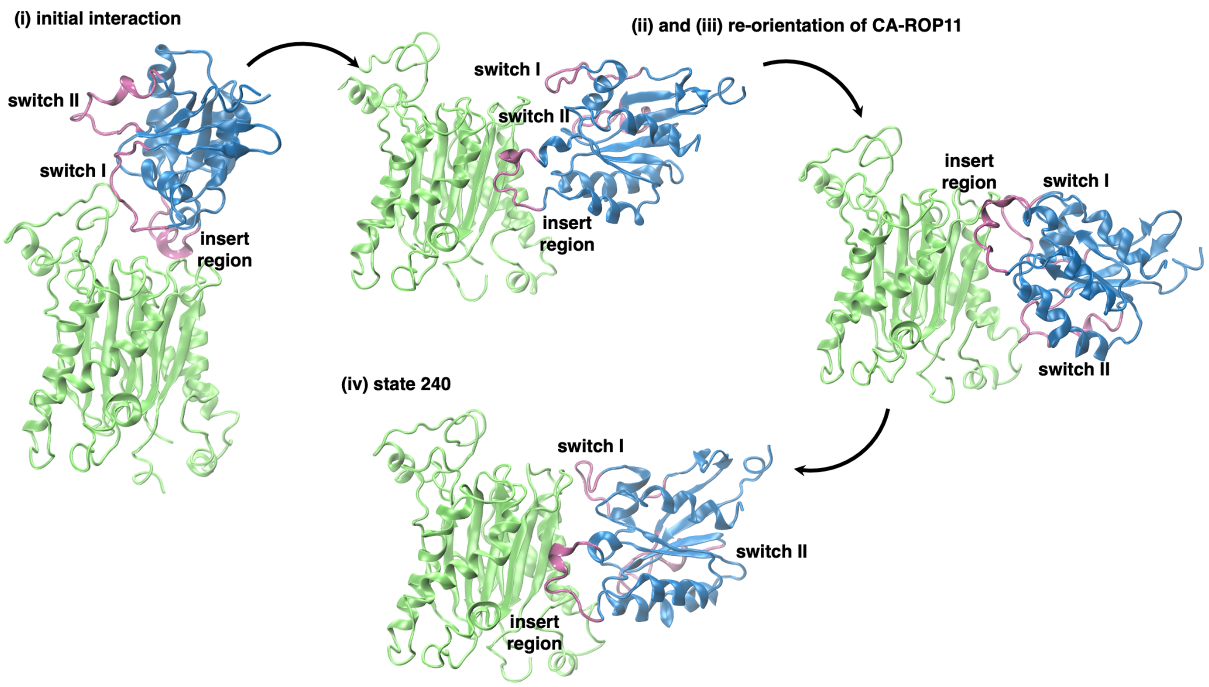


Figure S12: The association pathway between ABI1 and CA-ROP11 to form state 240 predicted from CG MD simulations. The initial interaction between ABI1 and CA-ROP11 is mediated by the charged residues in ABI1 and the insert region of CA-ROP11. Next, CA-ROP11 adapts the orientation to rotate around ABI1 and assumes the pose in state 240. This transition pathway was generated by performing transition path theory analysis on the MSM from CG MD simulations.

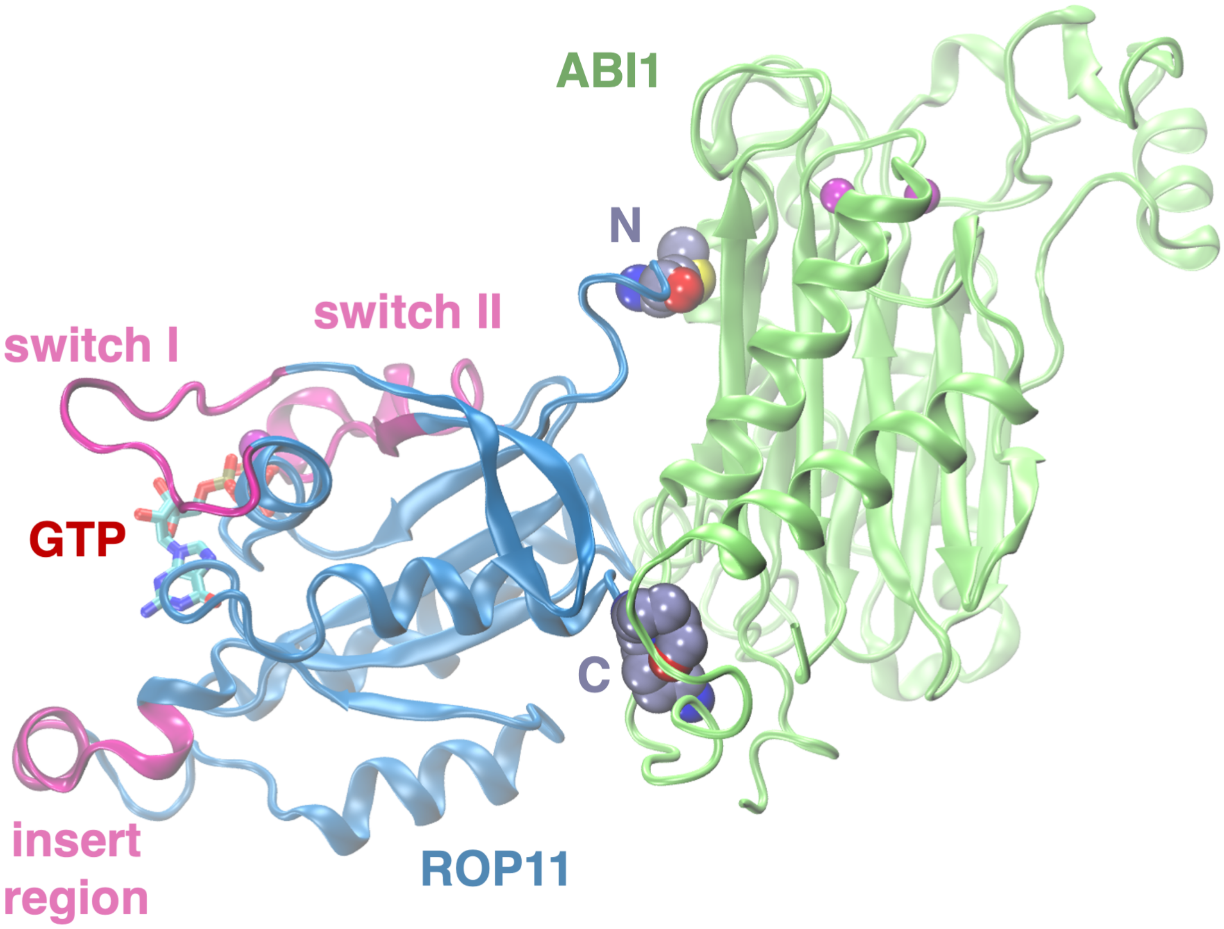


Figure S13: The candidate complex structure of the state 3 where both N- and C-terminal residues of ROP11 are involved in forming the interface, which is unlikely to be physical.

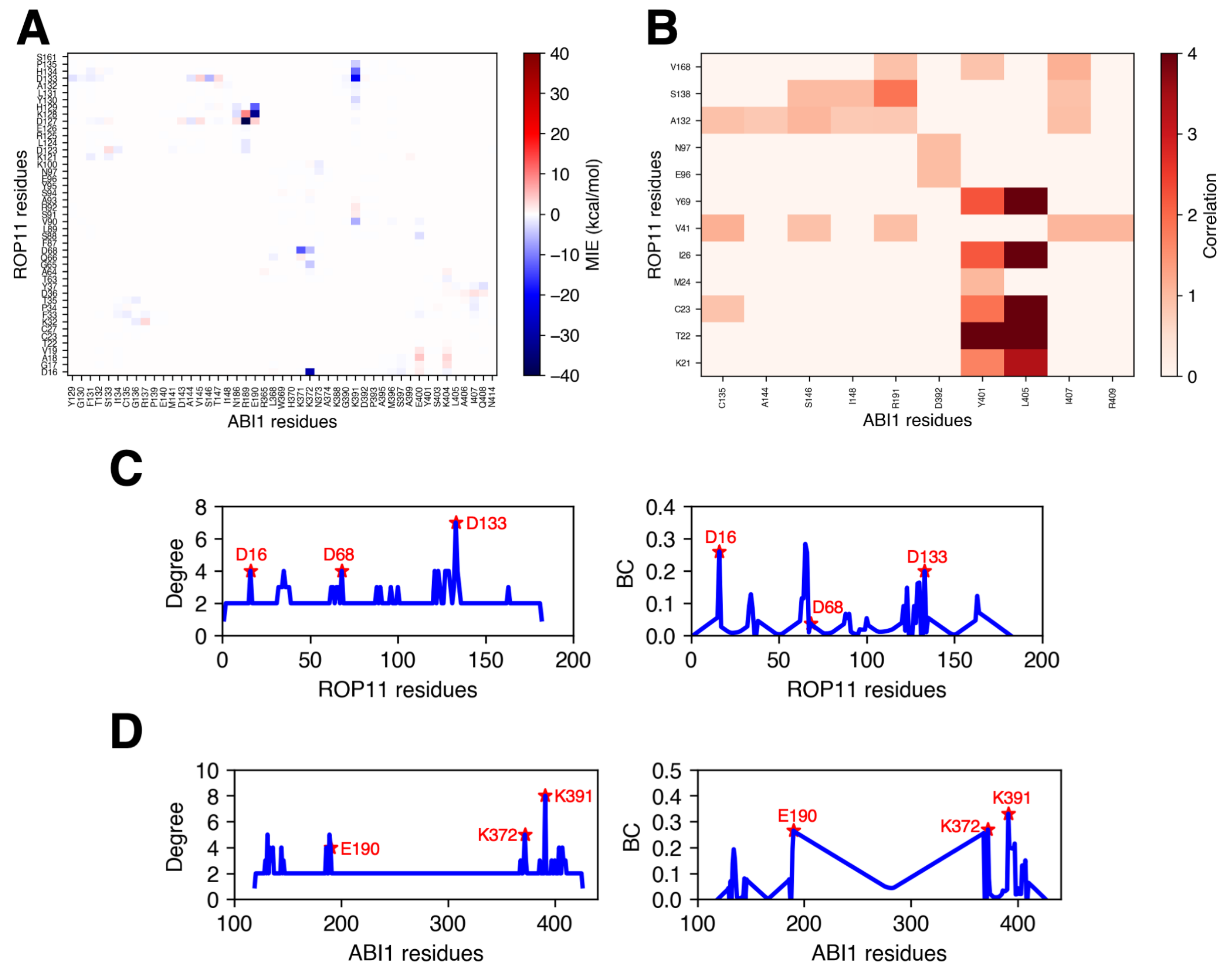


Figure S14: Identification of critical residue-residue pairs in state 240. (A) Mean interaction energy (MIE) matrix and (B) residue correlation matrix for the residue-residue pairs between ABI1 and ROP11. Network analysis reveals the degree and betweenness-centrality (BC) of the residues in (C) ROP11 and (D) ABI1. The residues in ROP11 and ABI1 with high degree and BC are highlighted.

(S4) Case, D. A.; Ben-Shalom, I. Y.; Brozell, S. R.; Cerutti, D. S.; Cheatham, T. E. I.; Cruzeiro, V.; Darden, T.; Duke, R.; Ghoreishi, D.; Gilson, M.; Gohlke, H.; Goetz, A.; Greene, D.; Harris, R.; Homeyer, N.; Izadi, S.; Kovalenko, A.; Kurtzman, T.; Lee, T.; LeGrand, S.; Li, P.; Lin, C.; Liu, J.; Luchko, T.; Luo, R.; Mermelstein, D.; Merz, K.; Miao, Y.; Monard, G.; Nguyen, C.; Nguyen, H.; Omelyan, I.; Onufriev, A.; Pan, F.; Qi, R.; Roe, D.; Roitberg, A.; Sagui, C.; Schott-Verdugo, S.; Shen, J.; Simmerling, C.; Smith, J.; Salomon-Ferrer, R.; Swails, J.; Walker, R.; Wang, J.; Wei, H.; Wolf, R.; Wu, X.; Xiao, L.; York, D.; Kollman, P. *Amber 18*; University of California: San Francisco, CA, 2018.

(S5) Loncharich, R. J.; Brooks, B. R.; Pastor, R. W. Langevin dynamics of peptides: The frictional dependence of isomerization rates of N-acetylalanyl-N'-

methylamide. *Biopolymers* **1992**, *32 . \aftergroup\futurelet\@let@token*, 523–535, doi:10.1002/bip.360320508.

(S6) Darden, T.; York, D.; Pedersen, L. Particle mesh Ewald: An Nlog(N) method for Ewald sums in large systems. *Journal of Chemical Physics* **1993**, *98*, 10089–10092.

(S7) Ryckaert, J.-P.; Ciccotti, G.; Berendsen, H. J. Numerical integration of the cartesian equations of motion of a system with constraints: molecular dynamics of n-alkanes. *Journal of Computational Physics* **1977**, *23*, 327–341.

(S8) Marrink, S. J.; Risselada, H. J.; Yefimov, S.; Tieleman, D. P.; de Vries, A. H. The MARTINI Force Field: Coarse Grained Model for Biomolecular Simulations. *The Journal of Physical Chemistry B* **2007**, *111*, 7812–7824.

(S9) Monticelli, L.; Kandasamy, S. K.; Periole, X.; Larson, R. G.; Tieleman, D. P.; Marrink, S.-J. The MARTINI Coarse-Grained Force Field: Extension to Proteins. *Journal of Chemical Theory and Computation* **2008**, *4*, 819–834.

(S10) Li, Z.; Kang, J.; Sui, N.; Liu, D. ROP11 GTPase is a Negative Regulator of Multiple ABA Responses in Arabidopsis. *Journal of Integrative Plant Biology* **2012**, *54*, 169–179.

(S11) Li, Z.; Li, Z.; Gao, X.; Chinnusamy, V.; Bressan, R.; Wang, Z.-X.; Zhu, J.-K.; Wu, J.-W.; Liu, D. ROP11 GTPase Negatively Regulates ABA Signaling by Protecting ABI1 Phosphatase Activity from Inhibition by the ABA Receptor RCAR1/PYL9 in Arabidopsis. *Journal of Integrative Plant Biology* **2012**, *54*, 180–188.

(S12) Periole, X.; Cavalli, M.; Marrink, S.-J.; Ceruso, M. A. Combining an Elastic Network With a Coarse-Grained Molecular Force Field: Structure, Dynamics, and Intermolecular Recognition. *Journal of Chemical Theory and Computation* **2009**, *5*, 2531–2543.

(S13) Abraham, M. J.; Murtola, T.; Schulz, R.; Páll, S.; Smith, J. C.; Hess, B.; Lindahl, E. GROMACS: High performance molecular simulations through multi-level parallelism from laptops to supercomputers. *SoftwareX* **2015**, *1-2*, 19–25.

(S14) Bowman, G. R.; Ensign, D. L.; Pande, V. S. Enhanced Modeling via Network Theory: Adaptive Sampling of Markov State Models. *Journal of Chemical Theory and Computation* **2010**, *6*, 787–794.

(S15) Shukla, D.; Hernández, C. X.; Weber, J. K.; Pande, V. S. Markov State Models Provide Insights into Dynamic Modulation of Protein Function. *Accounts of Chemical Research* **2015**, *48*, 414–422.

(S16) Husic, B. E.; Pande, V. S. Markov State Models: From an Art to a Science. *Journal of the American Chemical Society* **2018**, *140*, 2386–2396.

(S17) Naritomi, Y.; Fuchigami, S. Slow dynamics in protein fluctuations revealed by time-structure based independent component analysis: The case of domain motions. *Journal of Chemical Physics* **2011**, *134*, 065101.

(S18) McGibbon, R. T.; Pande, V. S. Variational cross-validation of slow dynamical modes in molecular kinetics. *Journal of Chemical Physics* **2015**, *142*, 124105.

(S19) Harrigan, M. P.; Sultan, M. M.; Hernández, C. X.; Husic, B. E.; Eastman, P.; Schwantes, C. R.; Beauchamp, K. A.; McGibbon, R. T.; Pande, V. S. MSMBuilder: statistical models for biomolecular dynamics. *Biophysical Journal* **2017**, *112*, 10–15.

(S20) McGibbon, R. T.; Hernández, C. X.; Harrigan, M. P.; Kearnes, S.; Sultan, M. M.; Jastrzebski, S.; Husic, B. E.; Pande, V. S. Osprey: Hyperparameter Optimization for Machine Learning. *Journal of Open Source Software* **2016**, *1*, 00034.

(S21) Wassenaar, T. A.; Pluhackova, K.; Böckmann, R. A.; Marrink, S. J.; Tieleman, D. P. Going Backward: A Flexible Geometric Approach to Reverse Transformation from

Coarse Grained to Atomistic Models. *Journal of Chemical Theory and Computation* **2014**, *10*, 676–690.

(S22) Sugita, Y.; Kitao, A.; Okamoto, Y. Multidimensional replica-exchange method for free-energy calculations. *Journal of Chemical Physics* **2000**, *113*, 6042–6051.

(S23) Phillips, J. C.; Braun, R.; Wang, W.; Gumbart, J.; Tajkhorshid, E.; Villa, E.; Chipot, C.; Skeel, R. D.; Kale, L.; Schulten, K. Scalable molecular dynamics with NAMD. *Journal of Computational Chemistry* **2005**, *26*, 1781–1802.

(S24) Shirts, M. R.; Chodera, J. D. Statistically optimal analysis of samples from multiple equilibrium states. *The Journal of Chemical Physics* **2008**, *129*, 124105.

(S25) Woo, H.-J.; Roux, B. Calculation of absolute protein-ligand binding free energy from computer simulations. *Proceedings of the National Academy of Sciences* **2005**, *102*, 6825–6830.

(S26) Gumbart, J. C.; Roux, B.; Chipot, C. Standard Binding Free Energies from Computer Simulations: What Is the Best Strategy? *Journal of Chemical Theory and Computation* **2012**, *9*, 794–802.

(S27) Gumbart, J. C.; Roux, B.; Chipot, C. Efficient Determination of Protein–Protein Standard Binding Free Energies from First Principles. *Journal of Chemical Theory and Computation* **2013**, *9*, 3789–3798.

(S28) Zhao, C.; Shukla, D. Molecular basis of the activation and dissociation of dimeric PYL2 receptor in abscisic acid signaling. *Physical Chemistry Chemical Physics* **2022**, *24*, 724–734.

(S29) Serçinoğlu, O.; Ozbek, P. gRINN. *a tool for calculation of residue interaction energies and protein energy network analysis of molecular dynamics simulations* **2018**, Nucleic Acids Research *46*, W554–W562.
